# Bridging scales in scattering tissues via multifocal two-photon microscopy

**DOI:** 10.1101/2020.06.11.146704

**Authors:** David Chen, Fabian Segovia-Miranda, Noreen Walker, Jose I. Valenzuela, Marino Zerial, Eugene W. Myers

**Author notes:** **Correspondence** Correspondence and requests for materials should be addressed to D.C.

## Abstract

Imaging biological systems at subcellular resolution and across scales is essential to understanding how cells form tissues, organs, and organisms. However, existing large-scale optical techniques often require harsh tissue-clearing methods that cause significant morphological changes, compromise the integrity of cell membranes, and reduce the signal of fluorescent proteins. Here, we demonstrate multifocal two-photon microscopy that enables imaging mesoscopic scattering samples in their native tissue environment at high resolution and high speed.

---

How cells determine the diversity of tissues and how they affect the state of health of organs and organisms are outstanding questions in biology. To understand the three-dimensional (3D) organization of cell types conforming organs and networks^1^, we must image tissues not only at high spatial resolution but also at large scales, as the principles governing the complexity of cellular structures cannot be elucidated from small-volume measurements.

Recent advances in light-sheet microscopy and tissue clearing have enabled whole-organ and whole-body imaging across scales^2, 3^. Unfortunately, light-sheet microscopy is prone to photon crosstalk that severely degrades the image contrast in strongly scattering tissues^4^ (for example, in brain, spleen, and liver) and its optical resolution in mesoscopic samples is typically of a few microns^3^, which is not sufficient for resolving subcellular structures accurately. Furthermore, large samples must be treated with harsh tissue-clearing agents, such as organic solvents, that often bleach fluorescent proteins, affect membrane integrity, and alter morphometric parameters^5–7^— essential for quantitative analysis and modeling of tissue function^8^. Therefore, multiscale microscopy requires techniques capable of sampling mesoscopic tissues at high resolution and great penetration depth without relying on aggressive clearing methods.

Two-photon microscopy has established itself as the gold standard for imaging optically challenging tissues at submicron resolution. Point-scanned detection is resistant to scattered photons and near-infrared wavelengths penetrate deeper into tissues than visible light^9^. However, large-scale measurements have been greatly hindered by the low throughput of two-photon microscopy. Indeed, it takes more than a week for a state-of-the-art two-photon microscope to image an entire mouse organ with a single fluorescent label^10^ or many months in the case of samples with multiple markers. Such long measurements are problematic given tissue degradation and loss of fluorescence signal in the sample over time^11^.

To bridge the gap between mesoscopic samples and high-resolution measurements, we have developed a multifocal scheme for two-photon microscopy that enables a 16-fold increase in acquisition speed compared with a conventional single-beam configuration. Different multifocal approaches have been demonstrated previously^12^. However, low image contrast due to inter-beam crosstalk has limited their practical applications^13–15^. By stabilizing the position of the multibeam fluorescence signal at the detector via emission re-scanning, we were able to probe deep tissues at high throughput without sacrificing the image quality. Furthermore, in combination with *in situ* tissue sectioning, we showed that multifocal scanning allowed imaging large volumes of optically challenging samples routinely. This cannot be achieved with conventional light-sheet or two-photon microscopy because of photon crosstalk in the former—even after tissue clearing^16^— and low acquisition throughput in the latter. We demonstrated the capabilities of this imaging platform by comprehensively mapping the cell nuclei and plasma membranes of an entire mouse liver lobe—several millimeters in size—in less than a day. The liver poses an exceptional challenge for microscopy because of its high tissue opacity, which has thus far hindered large-scale 3D measurements.

Our multifocal scheme performs fluorescence excitation by focusing an array of 16 collinear laser beams onto the sample (Fig. 1). The 16 laser foci divide the field of view (FOV) into equal-sized strips that are raster-scanned simultaneously in the fast and slow axes, *x* and *y*, respectively (Fig. 1.i). Epifluorescence is detected through a 16-channel photomultiplier tube (PMT) with the anodes linearly arranged along the *y*-axis (Fig. 1.ii). Our key innovation lies in re-scanning the multibeam fluorescence emission onto the PMT with a galvanometer mirror to keep the signal stationary along the *y*-axis but mobile within the anodes in *x* (Fig. 1.ii). Consequently, the signal crosstalk between the channels of the PMT and the optical loss in the inter-anode gaps are both minimized. We refer to this scheme as two-photon microscopy with **co**llinearly **m**ultiplexed **b**eams (2pCOMB).

**Fig. 1:**
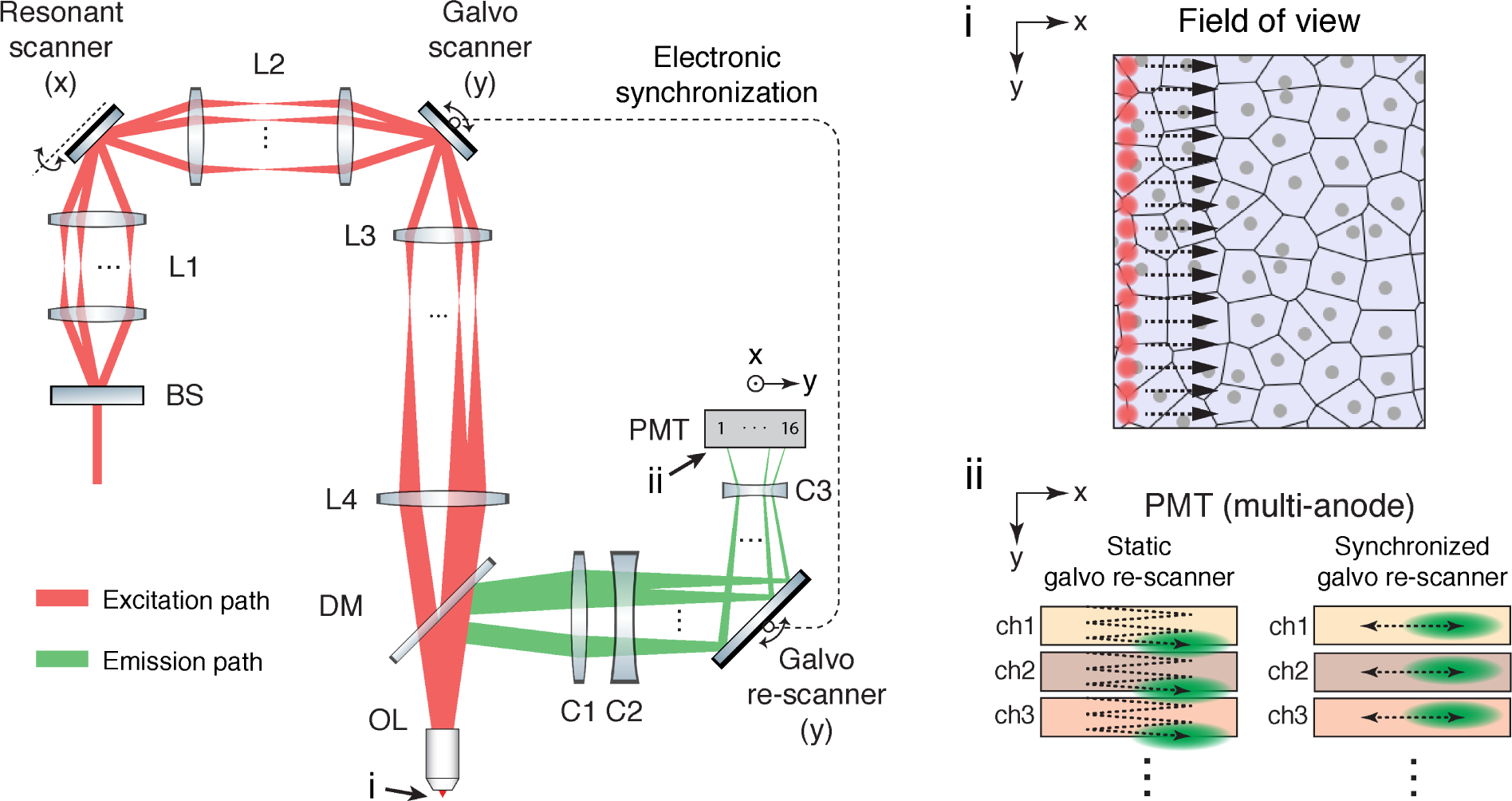
Simplified optical layout of two-photon microscopy with collinearly multiplexed beams (2pCOMB). A beam-splitter (BS) creates a linear array of 16 laser beams. The beams pass through relay lenses (L1–L4), focus onto the sample through an objective lens (OL) and scan the field of view with a ‘laser comb’ of 16 excitation foci (i). This beam configuration allows a 16-fold increase in throughput compared with a conventional single-beam scheme. The 16 fluorescence signals return through OL, reflect off a dichroic mirror (DM), and focus onto a 16-anode PMT through collector lenses (C1–C3). A galvanometer mirror in the *y*-axis re-scans the fluorescence emission to minimize the crosstalk between adjacent anodes of the PMT (ii). A detailed diagram can be found in Supplementary Fig. 1.

By imaging fluorescent beads via 2pCOMB (Fig. 2.a), we observed that the 16 parallel beams scanned the FOV simultaneously with minimal crosstalk. Residual crosstalk is induced by the nonzero spot size of the fluorescence beams focusing onto the anodes of the PMT as well as crosstalk between the emission foci at the sample plane due to scattered photons^9^. Additionally, emission from a uniform fluorescent slide showed a high signal to noise ratio across the FOV (Fig. 2.b) despite the uneven laser illumination of the diffractive beam splitter (Online Methods). These imaging artifacts can be effectively corrected through postprocessing, which is particularly straightforward in 2pCOMB because of the linear arrangement of the fluorescence beams. To this end, we developed methods for reducing the crosstalk and uneven illumination in the images (On-line Methods), which we evaluated on biological tissue. We used an established transgenic-mouse line expressing plasma-membrane-targeted tdTomato^17^ and stained with the dye DAPI that labels the double-stranded DNA in the cell nuclei (Online Methods). We compared postprocessed images scanned via 2pCOMB against control measurements acquired with the same microscope under a conventional single-beam configuration (Fig. 2.d–f and Supplementary Video 1). Comparable results between multifocal and single-beam imaging, both in shallow and deep tissues, demonstrate that 2pCOMB enables rapid two-photon microscopy without sacrificing the image contrast, resolution, or penetration depth.

**Fig. 2:**
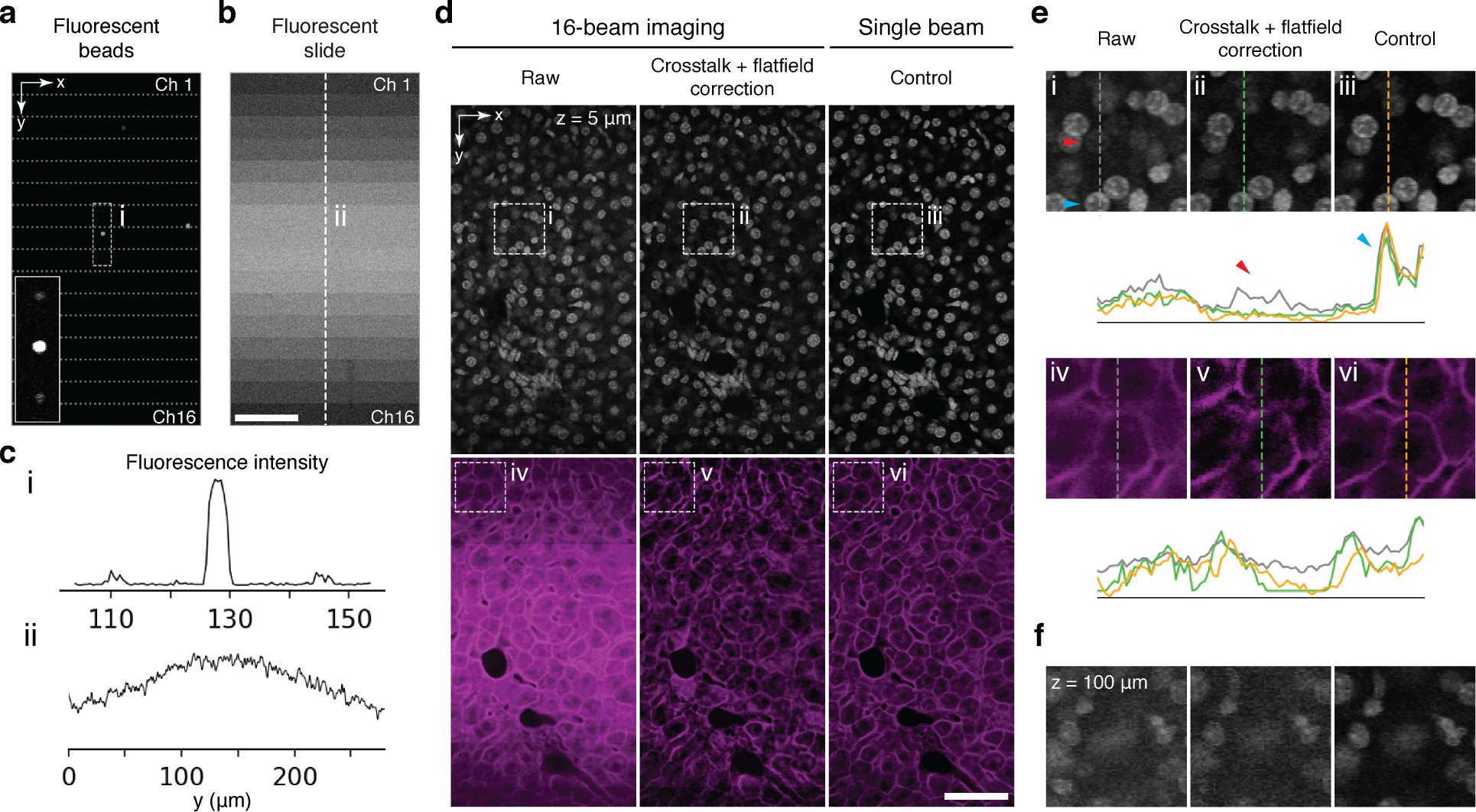
Fast and deep imaging with 2pCOMB. **a,** Fluorescent beads acquired via 2pCOMB. The FOV (150 µm × 280 µm) consists of 16 vertically-concatenated strips that are scanned simultaneously. Each strip contains the fluorescence signal of each anode of the PMT. The inset has been saturated to show the residual crosstalk between adjacent strips. **b,** Fluorescence emission from a uniform test slide acquired via 2pCOMB. **c,** Intensity profiles of a and b. **d,** Comparison of mouse liver tissue imaged with and without beam multiplexing for the cell nuclei (top row) and the plasma membranes (bottom row). Imaging artifacts have been reduced through postprocessing. The brightness of the images has been equalized to facilitate cross-comparison. **e,** Enlarged views of the selected areas in d and the corresponding intensity profiles. Red arrows point to signal crosstalk. **f,** Liver nuclei imaged at a depth of 100 µm (more details in Supplementary Note 9). All the images have been frame averaged 10 times and unwarped in *x* to correct for resonant scanning. Pixel size, 0.5 µm × 0.5 µm. Scale bars, 50 µm.

We next demonstrated that 2pCOMB allowed rapid volumetric imaging of large samples, including optically challenging specimens that cannot be adequately probed with light-sheet microscopy^16^. To this end, we imaged an entire caudal lobe of fixed mouse liver (~5 mm × 7 mm × 3 mm) under the same experimental conditions as in the previous measurement. Liver tissue is particularly challenging to probe due to its high opacity. Indeed, we have measured a two-photon penetration depth (at 1*/e*) of only ~46 µm in our samples, which severely hinders large-volume measurements (Supplementary Note. 6). High-resolution imaging of liver tissue has thus far been restricted to small volumes^8^ that do not capture the full structure of a single lobule (the basic functional unit of the lobe) or, even more challenging, the multi-lobule 3D organization of a whole lobe. To overcome the severe optical limitation in the liver that has hampered organ-scale measurements, we implemented *in situ* tissue sectioning^10^ to physically remove the surface of the sample after imaging the top layers at high speed via 2pCOMB (Fig. 3.a). Furthermore, for an accurate digital reconstruction of the lobe, we have increased the two-photon penetration in the liver to ~93 µm by means of the mild clearing protocol SeeDB^18^, which allowed us to image well below the layers of tissue damaged by the vibratome. SeeDB is an aqueous clearing method that makes the liver lobe semi-transparent without altering the membrane integrity, fluorescence of tdTomato^6^, or tissue morphology^7^ (Supplementary Notes 6–7).

**Fig. 3:**
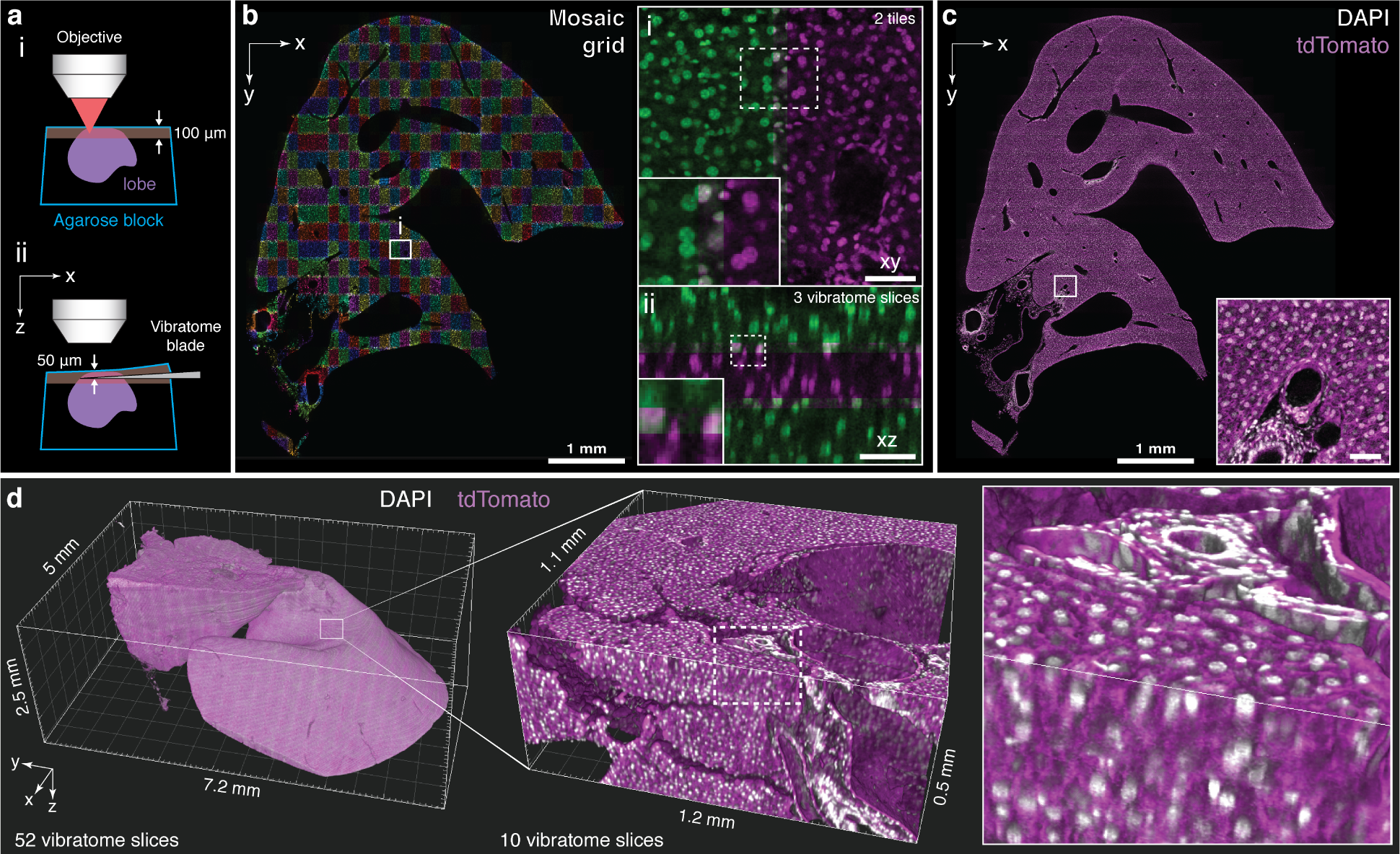
Multiscale imaging and digital reconstruction of a mouse liver lobe. **a,** An entire liver lobe was imaged iteratively by scanning its surface at high speed via 2pCOMB and then physically removing the tissue through *in situ* tissue sectioning (i and ii, respectively). **b,** Cell nuclei across a full *xy*-section of the lobe acquired via mosaic imaging with a *z*-stack of 150 µm × 280 µm × 100 µm in *xyz*. The enlarged views show co-aligned *z*-stacks in *xy* (i) and *xz* (ii) with an overlap of 10%, 15%, and 17% in *xyz*. Co-localized nuclei between adjacent stacks appear in grey. **c,** Cell nuclei (grey) across the same plane as in b with the stacks fused and overlaid with a second fluorescent channel showing the plasma membranes (magenta). The cross-section is located at approximately the center of the lobe. **d,** 3D rendering of the entire liver lobe comprised of 52 vibratome slices. The sample was imaged with a voxel size of 0.5 µm × 0.5 µm × 1 µm throughout the full volume. The lobe has been downsampled for display purposes (2 µm/voxel for the whole lobe and 1 µm/voxel for the enlarged volume). Scale bars, 50 µm, unless otherwise stated.

To image the entire liver lobe efficiently, we first previewed the location, shape, and extent of its surface via 2D panoramic scanning (Online Methods). The panorama was then overlaid with a 2D mosaic grid to mark the regions overlapping with the lobe for subsequent 3D scanning at high resolution. At every active grid position, we acquired a *z*-stack of 150 µm × 280 µm × 100 µm with a voxel size of 0.5 µm × 0.5 µm × 1 µm in *xyz*. The multifocal acquisition rate was 77 megavoxels per second (4.8 megavoxels per second per beam). The laser power was increased exponentially with depth to compensate for the signal attenuation in deep tissues (Supplementary Note 7). The initial power per beamlet was 30 mW at 1040 nm for the membranes and 5 mW at 750 nm for the nuclei. Between *z*-stacks, motorized stages repositioned the sample with submicron precision in a column-by-column manner with an overlap of 10% in *x* and 15% in *y* (Fig. 3.b). After imaging the surface of the lobe to a depth of 100 µm for both fluorescence channels, we physically removed the top 50 µm of the sample via *in situ* tissue sectioning to expose deeper layers (Fig. 3.a). The acquisition steps—panoramic previewing, mosaic imaging, and tissue sectioning—were then repeated until the entire volume was probed. Each vibratome slice had, therefore, a thickness of 100 µm and an image redundancy of 50% with the neighboring slices. This large overlap allowed us to reconstruct the lobe accurately despite the damage in the tissue caused by the vibratome.

For the entire liver lobe, we collected ~20,000 stacks for each of the fluorescence labels distributed over 52 vibratome slices. The voxel intensity was stored in 8 bits, resulting in a *z*-stack size of ~17 MB and a lobe size of ~330 GB per label. The acquisition time was ~11 hours per fluorescence label with a frame average of 2 to improve the SNR (Supplementary Table 2).

To digitally reconstruct the lobe, we first postprocessed all the *z*-stacks to correct for residual crosstalk, uneven illumination, field warping, and residual pixel-intensity variation in *z* (Online Methods). We then discarded the damaged and distorted layers of tissue induced by serial sectioning. Rigid stitching (Online Methods) was accurate for most of the lobe at the micron level (Fig. 3.b and Supplementary Videos 2–3). Only a few regions showed a mismatch of at most 2 nucleus diameters between vibratome slices (Supplementary Fig. 15), which could be improved via non-rigid stitching methods^19^.

A 3D rendering of the whole lobe (~41 mm^3^) at different magnification levels demonstrates the ability of our microscopy platform to image large volumes of tissue across multiple scales ranging from subcellular to quasi-organ-wide (Fig. 3.d and Supplementary Videos 4–8). Notably, we were able to image the 3D organization of the veins in the liver within and across the lobules, as well as the microscopic structures of the cell nuclei and plasma membranes throughout the entire lobe, which remained intact after optical clearing. This was possible even for a dense and opaque organ such as the liver, showing the capabilities of our imaging scheme.

The application of our microscopy platform to image an entire liver lobe rapidly and at high resolution demonstrates how 2pCOMB allows visualizing large volumes of tissue across multiple scales. We expect that this technology will enable the creation of comprehensive cell atlases and multiscale models, leveraging our systems-level understanding of tissue organization and development in health and disease.

## Data availability

Small datasets for the figures will be made available via the public repository of the MPI-CBG at https://git.mpi-cbg.de/. Large datasets for the whole liver lobe are available upon request.

## Code availability

All code used for analysis will be made available via the public repository of the MPI-CBG at https://git.mpi-cbg.de/.

## Acknowledgements

We would like to thank the Light Microscopy, Scientific Computing, Computer Department, International Office, and Scientific Infrastructure facilities of the MPI-CBG for their assistance; Nicola Maghelli (MPI-CBG) for reviewing and commenting on the manuscript; Martin Weigert (EPFL) for providing the code for unwarping the images. D.C. gratefully acknowledges support from the German Federal Ministry of Education and Research (BMBF), Grant no. 031L0044.

## Author contributions

D.C. and E.W.M. conceived the project. D.C. designed and built the microscope. D.C. wrote the control and acquisition software. F.S. established the clearing protocol for the liver. F.S and J.V. developed the staining methods and prepared the samples. D.C. conducted the experiments and postprocessed the images. N.W. digitally reconstructed the liver lobe and rendered the images and videos. D.C. lead and coordinated the project. M.Z. and E.W.M. acquired the financial support. D.C. wrote the paper with input from all co-authors.

## Competing Interests

The authors declare that they have no competing financial interests.

## Methods

### Custom two-photon microscope

We used a tunable laser (Coherent Vision II, 680–1080 nm, 140 fs, 80 MHz) and a fiber laser (Coherent Fidelity HP, 10 W, 1040 nm, 140 fs, 80 MHz) to excite blue and red fluorescence, respectively. The laser sources had enough laser power (*>*1 W) to drive 16 parallel excitation beams. The intensity of each laser was modulated with independent electro-op-tical modulators (Linos LM0202) and optical shutters (Uniblitz LS3S2Z1). Both laser beams were independently expanded and then combined with a polarizing cube (Semrock PBS1005-GVD). We used a diffractive optical element (DOE) for each laser source to split the light into an array of 16 beamlets (Holoor 16×1 array, 750 nm and 1040 nm). The DOEs were mounted in a motorized filter wheel (Thorlabs FW102C) for serial multicolor imaging. We used a resonant mirror (Cambridge Technology CRS 8 kHz) to scan the beam array in the fast axis *x* and a galvanometer mirror (Cambridge Technology 6215H, 5-mm mirror) for the slow axis *y*. The beams then passed through a Plössl scan lens (Thorlabs AC300-100-B), a tube lens (Thorlabs AC508-200-B-M), and an objective lens designed for two-photon imaging (Olympus XLSLPLN25XGMP, 25×, NA 1.0).

We mounted the sample on an acrylic container filled with immersion oil at room temperature. The container was attached to a tilt stage (Thorlabs PY004/M) and three motorized stages (Physik Instrumente; *x*: V-551.4B, *y*: V-551.2B, and *z*: ES-100) that positioned the sample in *xyz* with high precision. We used an *in situ* vibratome (Leica VT1200S) with stainless-steel blades (752-1-SS, Campden) to perform tissue sectioning.

### Re-scanned multifocal detection

The light emitted by the 16 fluorescence foci in the sample passed through the objective lens and reflected off a dichroic mirror (Semrock SEM-FF705-Di01-50-D) towards a 16-anode PMT (Hamamatsu H12311-40 MOD GaAsP). We designed an optical assembly of 2 collector lenses (Thorlabs AC254-100-A and ACN254-075-A) with the dual purpose of focusing the 16 fluorescence beams onto the detector plane and matching the separation of the beams to that of the anodes in *y*. We re-scanned the fluorescence beams onto the PMT with a galvanometer mirror (Cambridge Technology 6240H, 15-mm mirror) to minimize the crosstalk between the anodes in *y*. In contrast, the fluorescence spots were not re-scanned in *x*, and therefore, they could move inside the anodes as the sample was imaged across the fast axis. We prevented the fluorescence spots from moving beyond the sensing areas of the anodes by using a cylindrical lens (Newport CKX15-C, f = 25 mm) to reduce the optical magnification in this direction. We installed emission filters (Semrock FF01-492/sp and BLP01-532R) in a motorized filter wheel (Thorlabs FW102C) and a short-pass filter (Semrock F75-720) on the PMT to block stray light from the laser sources.

### Point spread function

We measured a PSF of 0.4 µm lateral and 2.6 µm axial at FWHM (Supplementary Note 5). The extraordinary long working distance (8 mm) of the selected objective, designed for imaging large and cleared whole samples, resulted in an axial resolution lower than that of objectives with similar characteristics but with a shorter WD. Nevertheless, imaging with in-situ tissue sectioning permits using an objective with a shorter WD to achieve a better axial resolution. Alternatively, isotropic resolution can be restored via postprocessing methods based on, for example, convolutional neural networks^1^.

### Control sequence and data acquisition

We wrote the main code in C++ and executed it on a desktop computer (Dell T7600, 64 GB RAM, 2 TB HDD, NVIDIA Quadro 4000, 10 G ethernet card). The computer interfaced with the microscope through an acquisition board with digital and analog inputs and outputs (National Instruments UBS-7856R). The board had an integrated FPGA processor that ran real-time routines programmable in LabView. The resonant scanner provided an electronic signal that served as a master clock for data acquisition. We synchronized both excitation and emission galvanometer scanners to minimize the inter-beam crosstalk between the anodes.

### Photocounting

For fast raster-scanning, we implemented an 8-kHz resonant mirror in the *x*-axis, which is typically used for *in vivo* imaging. We overcame the weak fluorescence signal inherent to rapid scanning by counting the number of single photons detected by the anodes (typically, 10–20 photons per pixel^2^). At low light conditions, photocounting presents a higher signal-to-noise ratio (SNR) than analog methods because of pulse-height discrimination^3^. In principle, the fluorescence signal could be improved by increasing the excitation power manyfold^4^. However, keeping the illumination intensity low prevents photobleaching the fluorescent markers—essential for serial multicolor imaging—and also permits supplying 16 two-photon excitation beams from a single power-limited light source. We built custom 16-channel electronic boards to discriminate the current pulses generated by the anodes of the PMT. We sampled the single-photon events during the pixel-dwell time with digital counters implemented on the FPGA. There were at most 13 counts per pixel for the particular dwell time of 162.5 ns and the laser repetition rate of 80 MHz used in the experiments (the pulse discriminator can only resolve one photon per laser pulse). Therefore, only 4 bits were needed for digitizing each pixel. The 16 independent counts were then multiplexed into two 32-bit numbers and sent to the computer through a FIFO queue. The computer demultiplexed the elements in the queue, upscaled each count from 4 bits to 8 bits, reconstructed the FOV, and then stored the image in a hard disk drive. The shot-noise in the images (Supplementary Fig. 10), common in measurements with low photocounts, can be effectively denoised through Gaussian filtering or deep-learning restoration^5^ (Supplementary Fig. 12), among other techniques.

### Panoramic previewing

After cutting the sample with the vibratome, we registered the location, extent, and shape of the new layers of tissue through 2 panoramic scans of ~10 mm × 15 mm at 30 µm and 90 µm below the surface. A panorama was formed by a collection of ribbon-shaped tiles (150 µm × 15 mm) stitched together in *x* to form a larger FOV (Supplementary Fig. 13). To acquire a ribbon-tile, the beam splitter in the microscope was bypassed to image under a conventional single-beam configuration. The resonant mirror scanned the single laser beam across the FOV in *x* while the sample was continuously translated with the *y*-stage^6^. We kept both excitation and emission galvanometer scanners fixed. We imaged the liver tissue via autofluorescence with the laser at 1040 nm and 15 mW at the focal plane. The acquisition time was ~1 minute per panorama with a pixel size of 0.5 µm × 1.0 µm. Panoramic previewing allows imaging samples of arbitrary shapes and sizes and is limited only by the travel range of the stages and the physical constraints of the sample mount.

### Volumetric *z*-stack imaging

For each *z*-stack, the 16 foci scanned the sample across 150 µm (300 pixels) bidirectionally in *x* at 8 kHz. In *y*, the laser foci traveled the inter-beam distance of 17.5 µm (35 pixels) at 29 Hz. The resulting multifocal acquisition throughput was 77 megavoxels per second (4.8 megavoxels per second per beam), which is equivalent to a frame rate of 460 Hz without frame averaging. In *z*, the vertical stage was continuously translated 100 µm (100 pixels), which led to a volume acquisition rate of 4.6 Hz (without frame averaging). We increased the laser power exponentially with depth to compensate for the signal attenuation in deep tissue. Each *z*-stack contained a volume of 150 × 280 × 100 µm^3^ in *xyz*. The voxel size was 0.5 µm × 0.5 µm × 1 µm in *xyz*, which had sub-Nyquist resolution in *z* and was sufficient to resolve the plasma membrane in *xy*. We used a stack overlap of 10% in *x* and a slightly larger value in *y*, 15%, to improve the uneven SNR in this direction due to non-uniform multifocal illumination.

### Vibratome slicing

For cutting the tissue, we set the vibratome to an amplitude of 1 mm and moved the sample towards the vibrating blade with the *y*-stage at 0.1 mm/s. Each cutting cycle was performed in ~1 minute.

### Fluorescence co-localization

We used the *xyz* motorized stages to correct for the chromatic shift in the excitation foci induced by aberration in the optical train and residual misalignment between the laser sources. For the liver, we visually co-aligned the cell nuclei and plasma membranes with sub-nuclear precision by examining the chromatic shift at the contour of the sample. The co-alignment remained stable throughout the whole-lobe acquisition. The focal shift between the blue and red emissions in our microscope was ~1 µm lateral and ~6 µm axial.

### Post-acquisition image processing

We applied two post-acquisition corrections to the images obtained via 2pCOMB. First, we reduced the residual crosstalk between the anodes of the PMT, detrimental to the image contrast, by subtracting the fluorescence signal from the adjacent anodes,
namely,

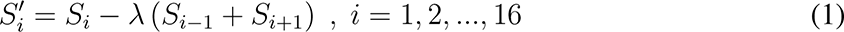

where *S_i_* is a strip of 300 × 35 pixels containing the fluorescence signal from the *i*-th anode and *λ* parametrizes the crosstalk correction. Second, we corrected the uneven laser illumination, induced by spatial smearing of the diffraction orders of the DOE^7^, by upscaling the pixel values strip-wise across the FOV (300 × 560 pixels) following the inverse-gaussian function

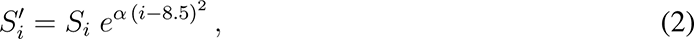

where *α* parametrizes the flatfield correction. Through quantitative analysis on our liver data, we found that the optimal parameters were *λ* = 0.28, *α* = .010 for the cell nuclei and *λ* = 0.35, *α* = 0.017 for the plasma membranes (Supplementary Note 8). Additionally, we unwarped all the images in the *x*-axis via standard pixel re-assignment to correct for the sinusoidal motion of the resonant scanner. The images presented in this paper have been postprocessed only with the aforementioned methods and do not involve denoising or deconvolution unless otherwise stated.

### Volume reconstruction via rigid stitching

We first corrected for residual intensity variation in *z* by normalizing the images with a reference curve obtained from averaging hundreds of *z*-stacks. Within each vibratome slice, we observed that the co-alignment of the *z*-stacks based purely on the stage position was accurate at the micron level (Fig. 3.b.i) and did not require any further image correlation. Therefore, we used the stage coordinates in the Fiji plugin Grid/Collection Stitching^8^ to stitch the stacks and fuse each vibratome slice independently (Fig. 3). We then computationally co-aligned the 52 vibratome slices in *xyz* (already individually fused) with the Fiji plugin BigStitcher^9^. We used the nuclei channel to compute the relative shifts between slices and then applied the result to both channels. For our specific liver sample, we found good alignment between neighboring vibratome slices, at the sub-nuclear level, after discarding the damaged and distorted tissue from serial sectioning enclosed within the top 40 µm of each *z*-stack (Fig. 3.b.ii) (the stack overlap in *z* decreased from 50% to 17%). In a small number of places, however, the tissue deformation induced by cutting penetrated deep inside the sample and resulted in a local misalignment of at most 2 nucleus diameters between neighboring vibratome slices (Supplementary Fig. 15). Future work may address such an effect through non-rigid transformations^10^. Finally, we rendered the whole lobe in BigStitcher and Imaris 9.5 (Fig. 3.d). The reconstruction was performed on a Dell T630 computer with 24 cores, 384 GB RAM, 144 TB HDD, NVIDIA Titan XP, 10 G ethernet card, and CentOS Linux. Additional details can be found in Supplementary Notes 13.

### Fluorescent slide and beads

For visualizing the field illumination and residual crosstalk, we used a plastic fluorescent slide (Chroma Technologies 92001) and a slide with 4 um fluorescent beads (FocalCheck #1, Fisher Scientific), respectively.

### Mouse liver samples, labeling, and optical clearing

We used the established transgenic mouse line C57BL/6J RosamT/mG^11^ that expresses membrane-targeted tdTomato to image the plasma membrane of the cells. Livers were fixed through transcardial perfusion with 4% paraformaldehyde, 0.1% Tween-20 in PBS, and post-fixed overnight at 4 °C with the same solution. Caudate lobes were dissected, stained, and optically cleared as previously described^12, 13^. Briefly, liver lobes were permeabilized with 0.2% gelatin, 0.5% Triton X-100 in PBS (PBSGT) for 4 days at room temperature (RT). We incubated the tissue with 2 µg/mL DAPI diluted in PBSGT with 0.1% 7 saponin for 14 days at 37 °C. Caudate lobes were washed overnight with PBSGT and embedded in 4% agarose in PBS. For optical clearing, we used a modified version of SeeDB^12^. Stained samples were consecutively incubated at RT in 25% wt/vol fructose for 4–8h, 50% wt/vol fructose for 4–8h, 75% wt/vol fructose for 16–20h, 100% wt/vol fructose for 48 h, and SeeDB (80.2% wt/vol fructose, 0.5% 1-thioglycerol, ~0.1M phosphate buffer pH7.5) for 48 h. Tissues were kept in SeeDB until imaging. As the immersion medium, we used a mix of 50% silicone and 50% mineral oil that matches the refractive index of SeeDB (1.49).

All procedures were performed in compliance with German animal welfare legislation and in pathogen-free conditions in the animal facility of the MPI-CBG, Dresden, Germany. Protocols were approved by the Institutional Animal Welfare Officer (Tierschutzbeauftragter) and all the necessary licenses were obtained from the regional Ethical Commission for Animal Experimentation of Dresden, Germany (Tierversuchskommission, Landesdirektion Dresden) (License number: DD24-5131/338/50).

## Supplementary Information

### 1 Abbreviations

2PM: Two-photon microscopy
DOE: Diffractive optical element
FOV: Field of view
PMT: Photomultiplier tube
MSE: Mean squared error
SNR: Signal-to-noise ratio
PSNR: Peak signal-to-noise ratio
2D: Two dimensional
3D: Three dimensional
PSF: Point spread function
WD: Working distance
PBS: Phosphate-buffered saline
RT: Room temperature
RI: Refractive index
SSIM: Structural similarity index

### 2 Detailed optical layout of 2pCOMB

**Supplementary Fig. 1:**
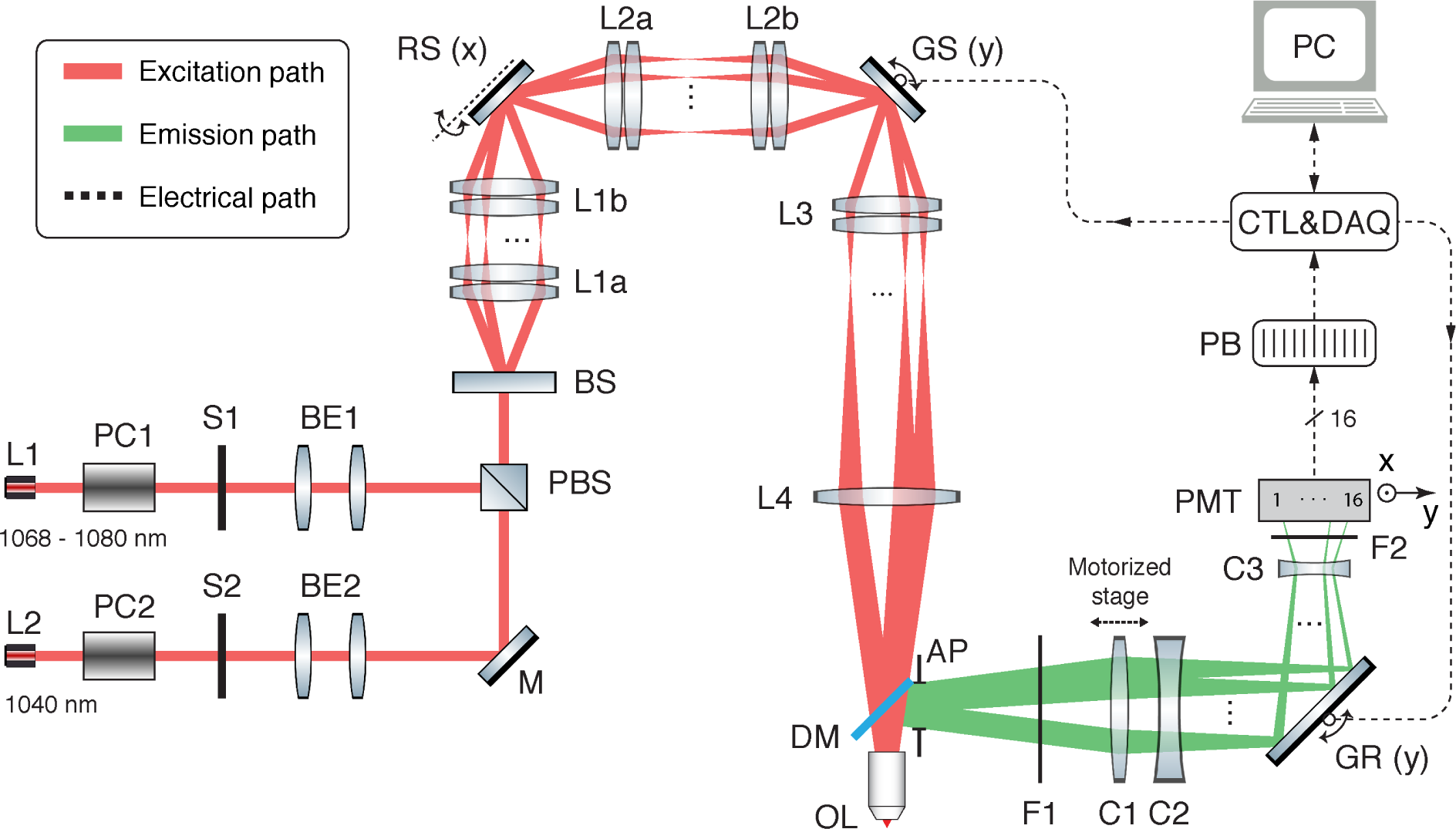
Detailed optical layout of 2pCOMB. Dimensions, lengths, and beam angles are not to scale. The beam angles were exaggerated for clarity. The full list of components can be found in Supplementary Table 1. Some optical symbols were obtained from ComponentLibrary^1^.

We adopted a combined achromat-meniscus lens for L1a, L1b, L2a, L2b, and two achromats in Plössl configuration for the scan lens L3 to minimize the optical aberration throughout the optical train^2^.

We found that the fluorescence emission produced by the laser source L1 (at 750 nm) was focusing on a slightly different plane than that of the laser L2 (at 1040 nm). This focal shift was caused by imperfect collimation of the laser sources L1 and L2 as well as chromatic aberration in the microscope. To be able to perform automated multicolor imaging, we mounted C1 on a motorized stage to refocus the fluorescence emission onto the PMT after switching lasers.

We found that the crosstalk between anodes is higher for fluorescence in the blue wavelengths than in the red. We attributed such an effect to an increase in the fluorescence spot size focusing onto the anodes of the PMT due to optical aberration, which is more prominent at short wavelengths. We were able to reduce the excess of crosstalk by installing an iris diaphragm (AP) in the detection path that served as an aperture stop (Supplementary Fig. 1).

### 3 List of electro-optical components

**Table 1:**
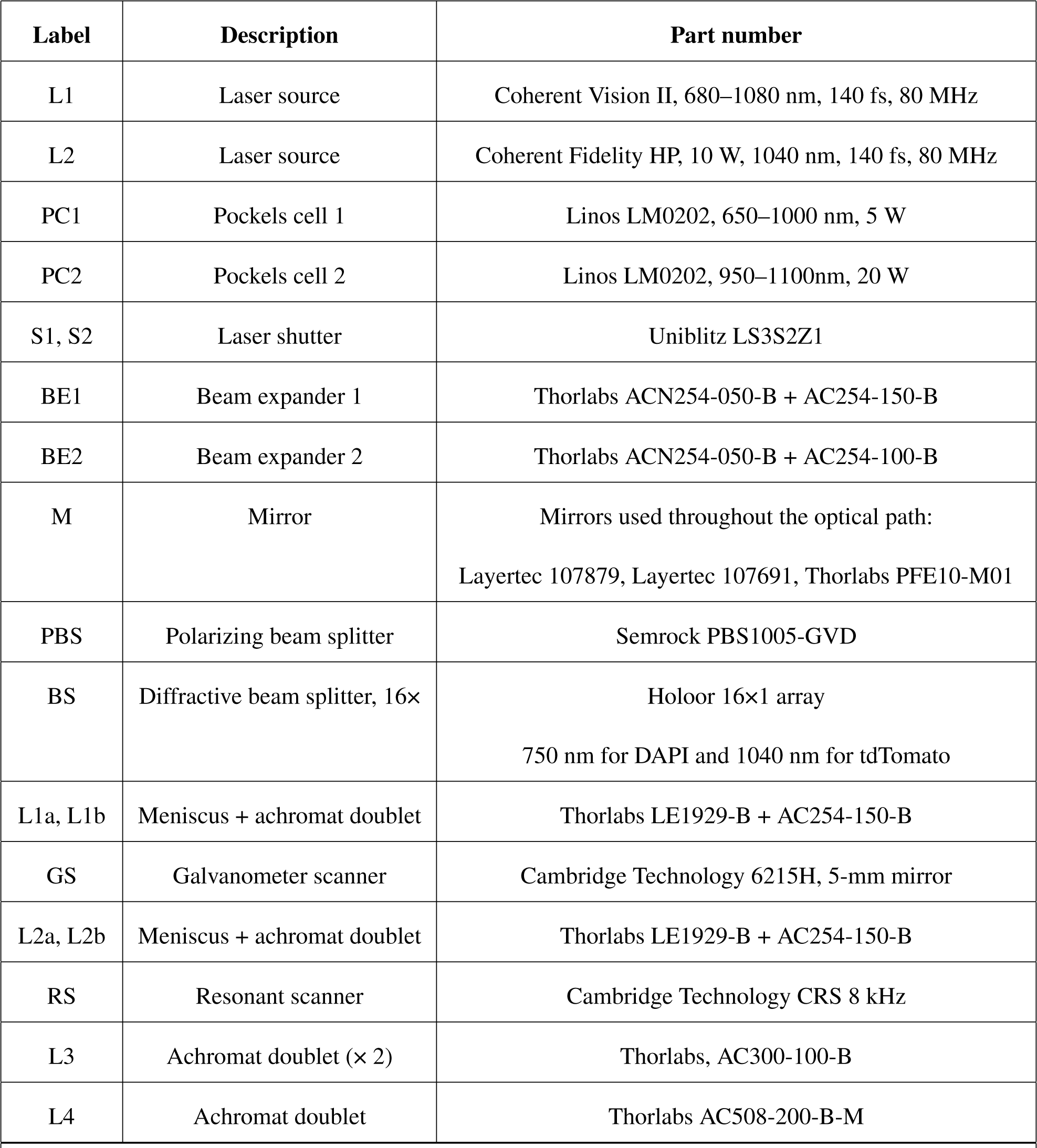
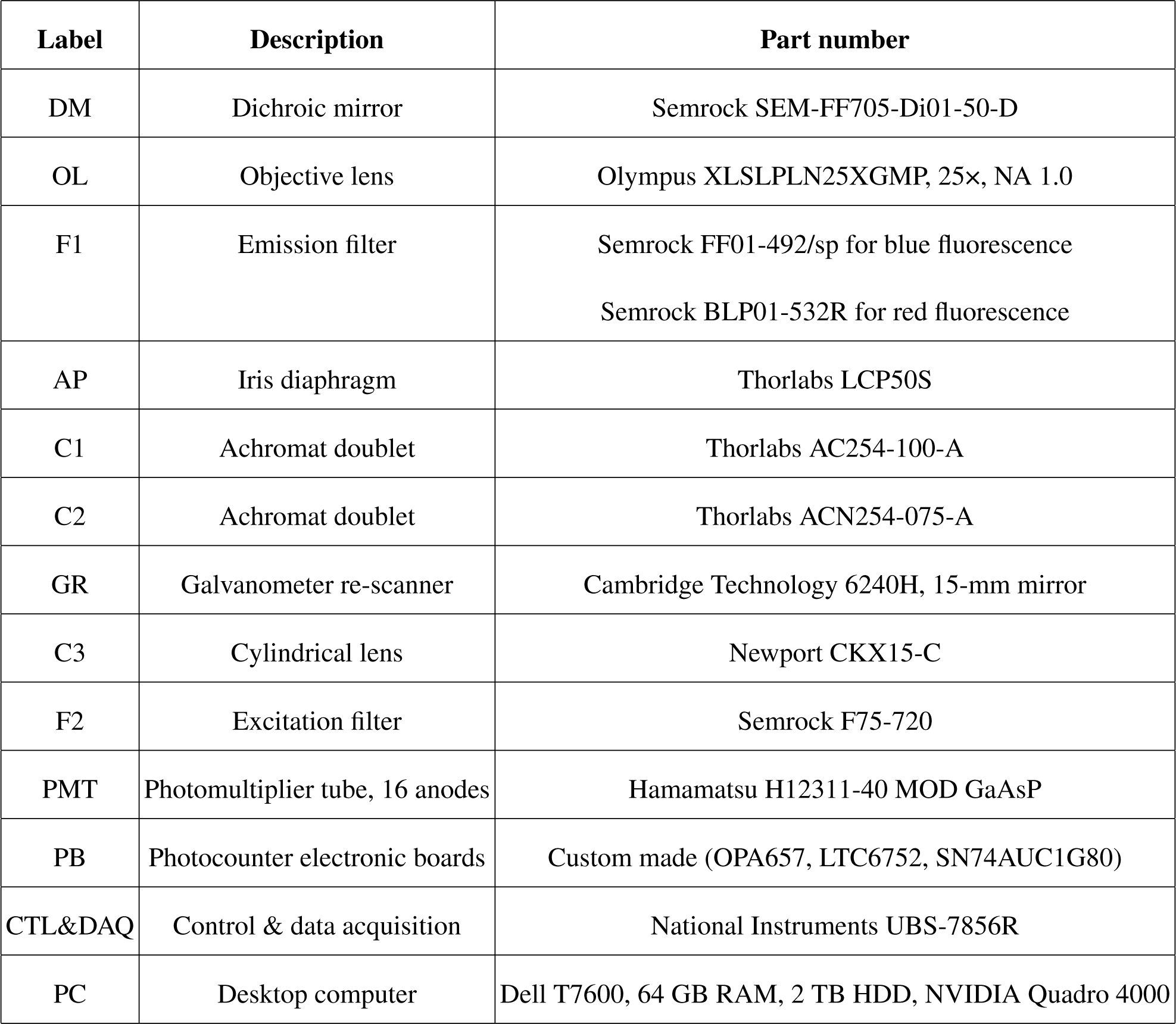
List of electro-optical components.

### 4 Three-dimensional optical layout of the fluorescence re-scanner in 2pCOMB

**Supplementary Fig. 2:**
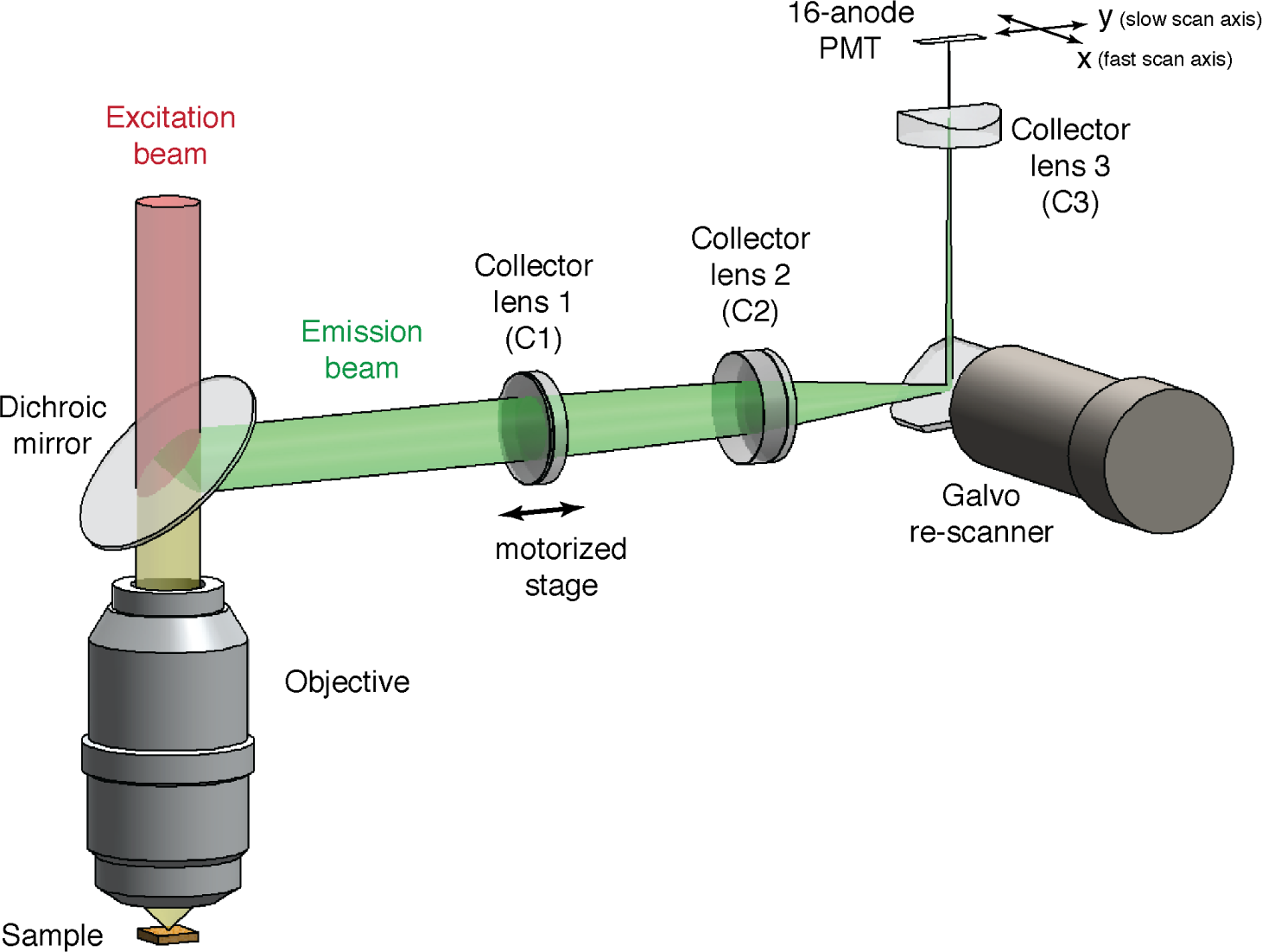
Three-dimensional optical layout of the fluorescence re-scanner in 2pCOMB. Dimensions, lengths, and angles are to scale. The emission filter, excitation filter, and iris diaphragm are not shown. Only a single beam in the laser comb is shown for clarity.

An optical assembly of 2 collector lenses (C1 and C2) was designed with the dual purpose of focusing the 16 fluorescence beams onto the anodes of the PMT in *y* and matching the inter-beam separation to that of the anodes, also in *y*. C2 is a cylindrical lens that loosely focused the beams onto the anodes in *x*. To allow automated multicolor imaging, we mounted C1 on a motorized stage to refocus the fluorescence emission onto the PMT after switching wavelengths (Supplementary Note 1).

### 5 Point spread function

To measure the point spread function (PSF) in our custom microscope, we used sub-diffraction fluorescent beads (Invitrogen TetraSpeck T7279, 0.1 µm, multicolor) mounted on a glass slide without a coverslip and immersed in silicone oil (RI, 1.51). For the particular objective lens used (Olympus XLSLPLN25XGMP, 25×, NA 1.0), we measured a PSF at 750 nm of 0.36 µm lateral and 2.59 µm axial at FWHM (Supplementary Fig. 3.a). This measurement was performed close to the upper edge of the FOV, and therefore, the bead was excited with the first beamlet of the laser array (see Fig. 7.a as a reference). The optical resolution in the rest of the FOV was, therefore, expected to be equal or better. The back aperture of the objective lens had a diameter of 14.4 mm and was filled with a gaussian laser beam with a diameter of ~14.4 mm at 1*/e*^2^. L1 was used as the excitation laser (Supplementary Fig. 1), which had a beam diameter of ~1.2 mm at the source and was expanded 3 times by BE1 and 4 times by the scan-tube lens assembly.

As a control measurement, we repeated the measurement with the microscope under a conventional single-beam configuration, this is, with the beam splitter bypassed. Under comparable experimental conditions, we obtained a PSF of 0.29 µm lateral and 2.23 µm axial at FWHM (Supplementary Fig. 3.b). We attributed the lower axial resolution of our microscope compared with similar measurements in the literature^3, 4^ to the extraordinary long working distance of our objective (8 mm) designed for imaging large and cleared whole samples. Nevertheless, imaging with in-situ tissue sectioning permits using an objective with a shorter WD to achieve a better axial resolution.

**Supplementary Fig. 3:**
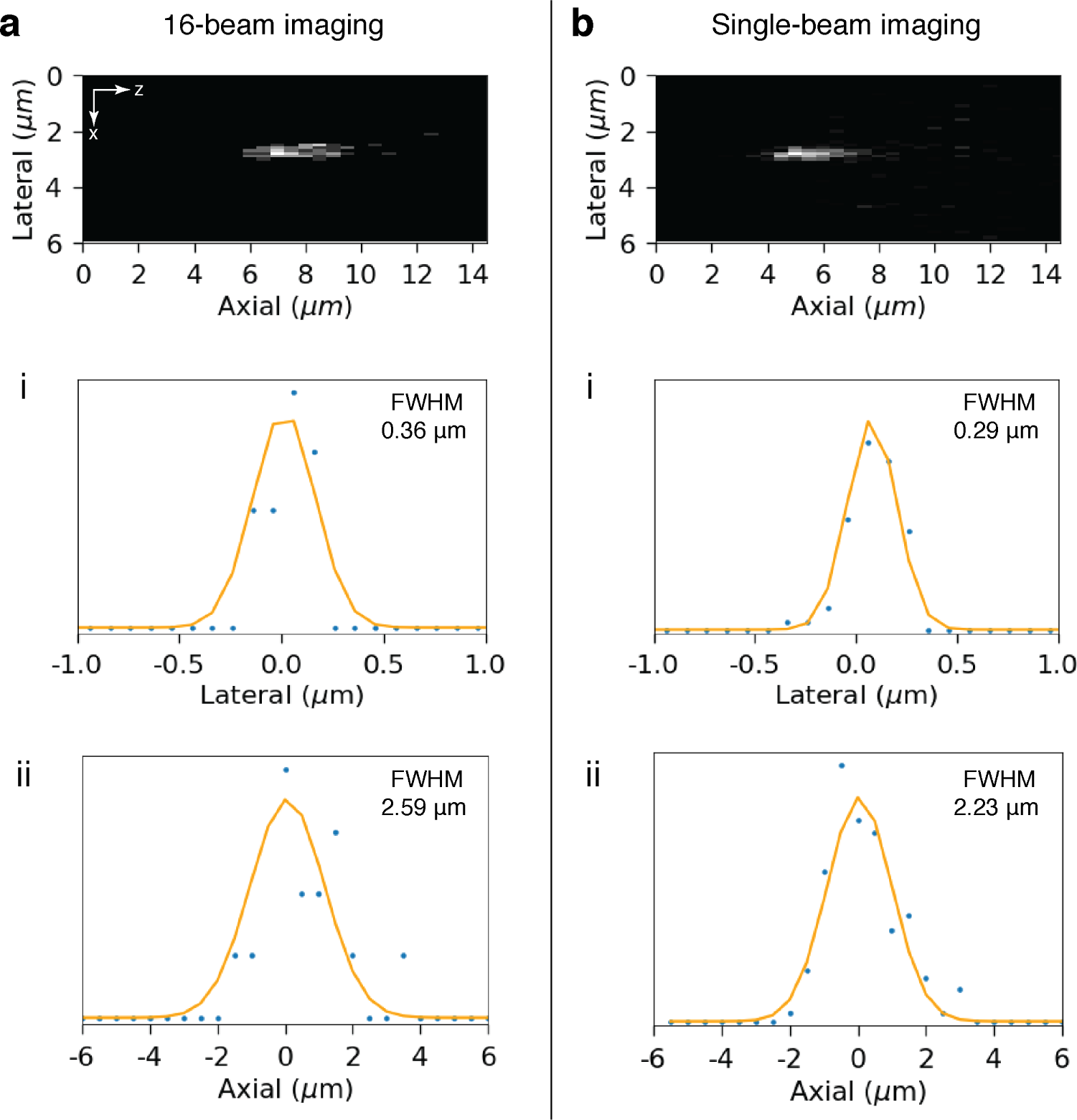
Point spread function. **a,** Fluorescent bead of diameter 0.1 µm in the *xz*-plane imaged via 2pCOMB at 750 nm. The bead was positioned at the upper edge of the FOV (see Fig. 7.a as a reference). Fluorescence emission was, therefore, excited with the first beamlet of the laser array. The bead was sampled volumetrically via *z*-scanning and fit with a 3D Gaussian function. The lateral and axial profiles are shown in i and ii, respectively. **b,** Control measurement acquired with the same microscope under a conventional single-beam configuration (i.e., with the beam splitter bypassed).

### 6 Imaging depth of two-photon microscopy in the mouse liver

We measured the imaging penetration depth of our custom two-photon microscope in the mouse liver before and after applying the tissue clearing protocol SeeDB^5^ (Online Methods). We imaged the plasma membranes and fit an exponential decay to the mean pixel intensity versus depth (Supplementary Fig. 4.c). The fit showed a decay length at 1*/e* of 46 µm ± 0.1 µm without tissue clearing and 93 µm ± 0.2 µm with SeeDB.

**Supplementary Fig. 4:**
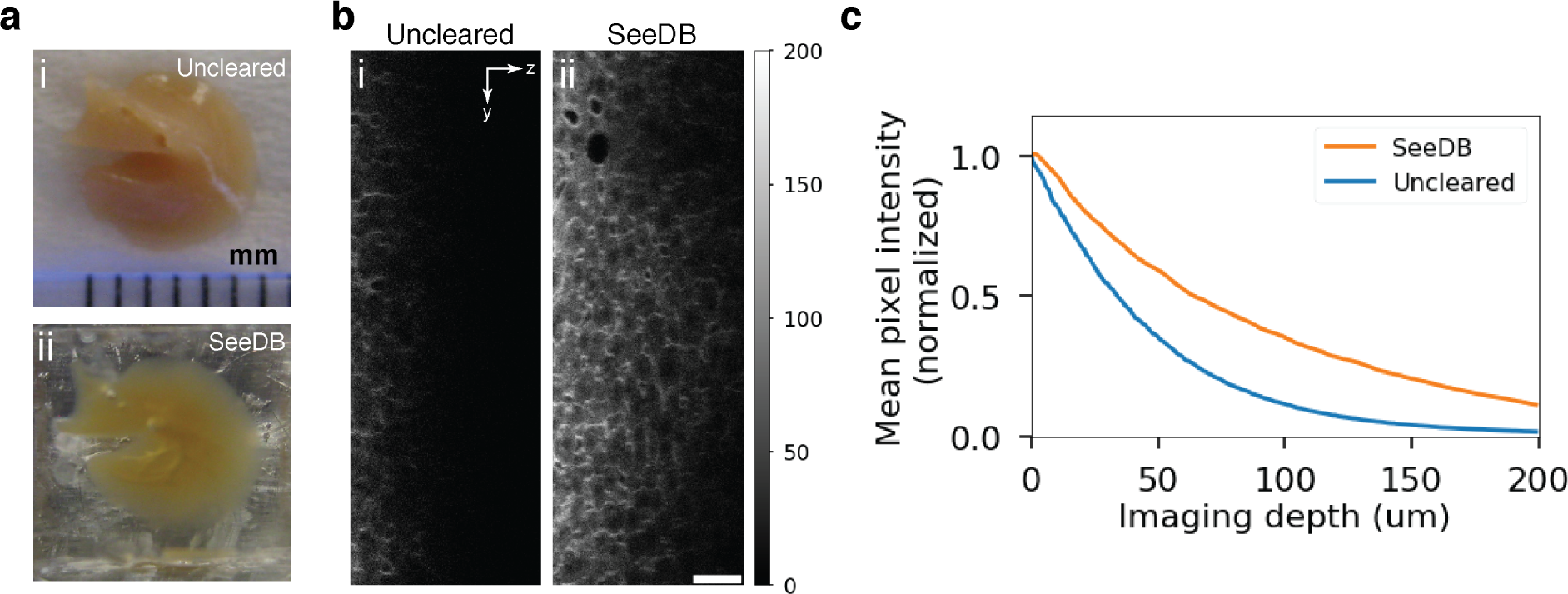
Two-photon imaging depth in the mouse liver. **a,** Photos of a caudal lobe before (i) and after (ii) tissue clearing with SeeDB. **b,** Plasma membranes across a *yz*-section with (i) and without (ii) tissue clearing. The images were acquired with our custom microscope under a conventional single-beam configuration (i.e., with the beam splitter bypassed). Pixel depth, 0–255. Scale bar, 50 µm. **c,** Mean pixel intensity (normalized) versus image depth with and without tissue clearing.

### 7 Depth-dependent laser power compensation

We compensated for the loss of fluorescence signal in deep tissues by increasing the two-photon laser power exponentially with depth. The resulting pixel intensity was approximately constant throughout *z* (Supplementary Fig. 5). Residual pixel intensity variation in *z* was reduced via post-acquisition processing (Online Methods and Supplementary Fig. 14).

**Supplementary Fig. 5:**
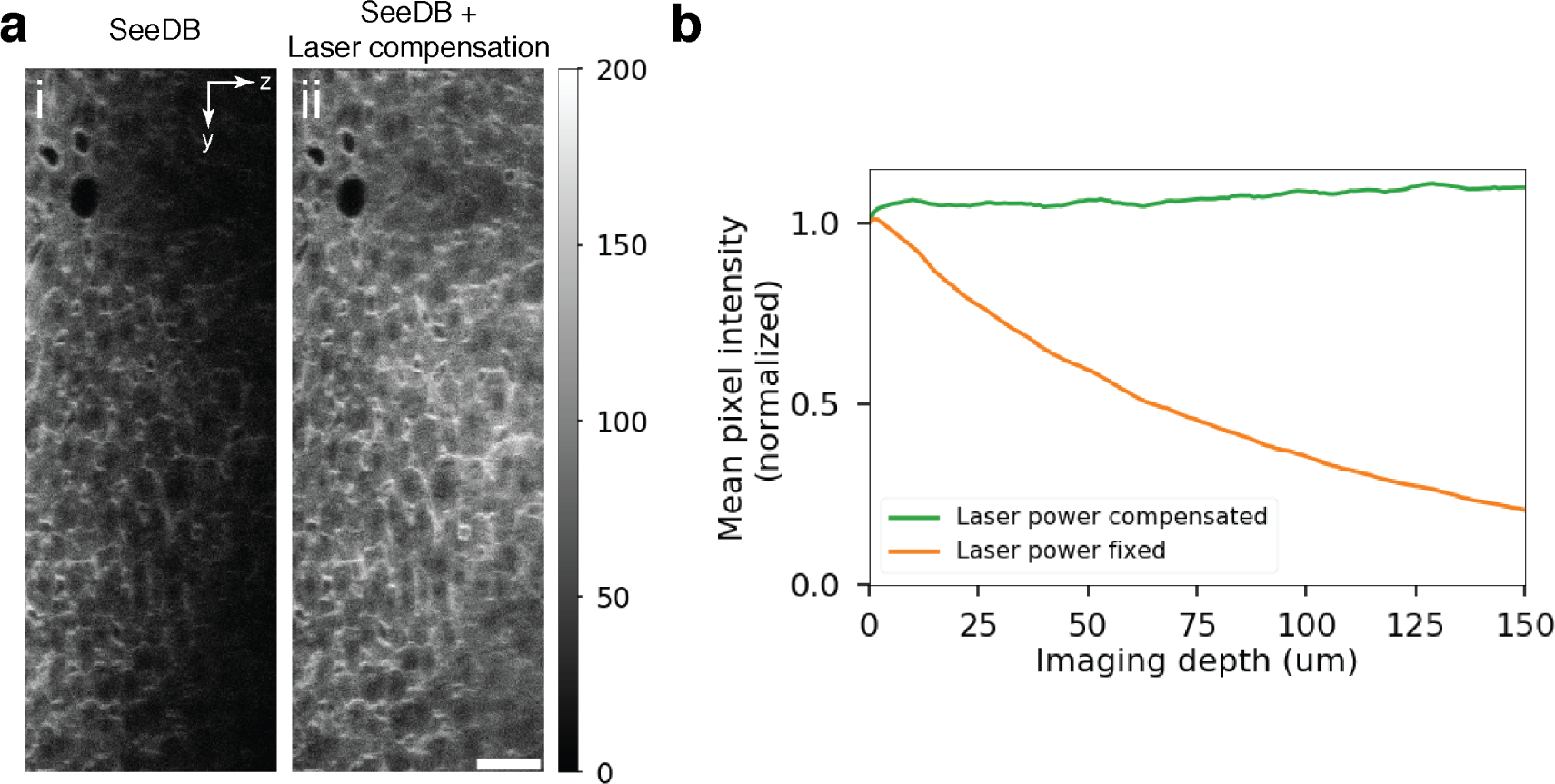
Imaging depth with laser power compensation. **a,** Plasma membranes across a *yz*-section without (i) and with (ii) laser power compensation in mouse liver tissue. The sample was cleared with SeeDB. The images were acquired with our custom microscope under a conventional single-beam configuration (i.e., with the beam splitter bypassed). Pixel depth, 0–255. Scale bar, 50 µm. **b,** Normalized mean pixel intensity versus image depth.

### 8 Post-acquisition processing

We reduce the uneven field illumination and residual channel crosstalk in 2pCOMB via post-acquisition processing, as described in Online Methods. A detailed visual summary of the postprocessing steps is shown in Supplementary Fig. 6. The images in Supplementary Fig. 6 correspond to those displayed in Fig. 2 in the main text. All the images in this paper have been additionally unwarped in *x* via pixel re-assignment with an inverse sine function to correct for the sinusoidal motion of the resonant scanner.

Improvements on 2pCOMB can potentially eliminate the need for post-acquisition image correction. For example, the stationary fluorescence spots along the *y*-axis of the PMT permits suppressing the inter-anode crosstalk entirely by mutually isolating the sensing areas with dividing walls. Furthermore, crosstalk between the fluorescent foci in the sample plane could be minimized via spatiotemporal multiplexing^6^. Finally, the uneven laser intensities of the beam splitter could be overcome by using instead a spatial light modulator optimized for field uniformity^7^.

**Supplementary Fig. 6:**
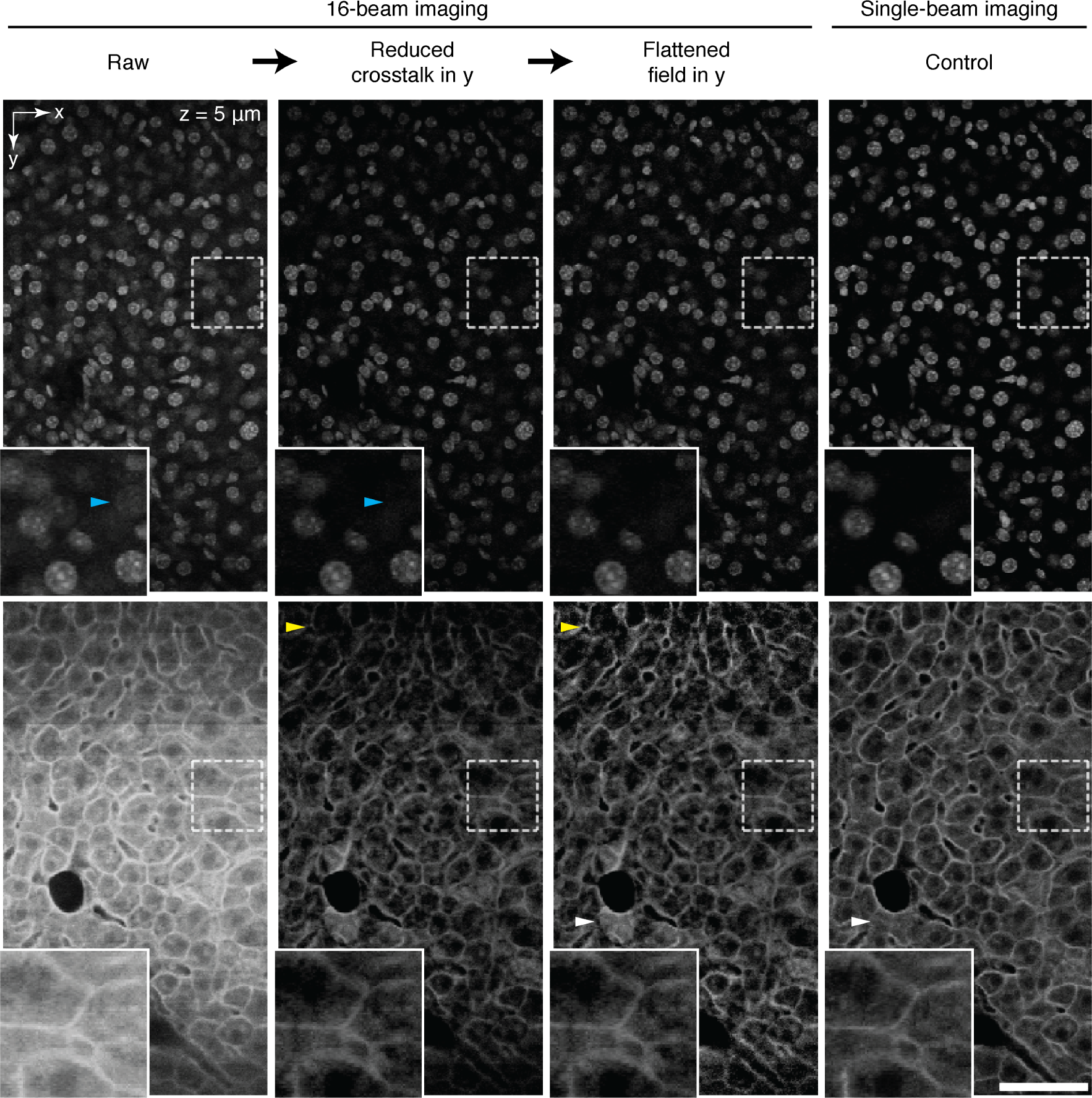
Summary of the postprocessing steps in 2pCOMB. Cell nuclei (top row) and plasma membranes (bottom row) in mouse liver tissue acquired with 16 parallel beams via 2pCOMB. The images show the postprocessing steps: field unwarping, crosstalk reduction, and flatfield correction. The control images on the right column were acquired with the same microscope under a single-beam configuration (i.e., with the beam splitter bypassed). The insets show enlarged views of the selected areas. The blue and yellow arrows point to particular areas with crosstalk and flatfield correction, respectively. The white arrows point to a postprocessing artifact resulting from applying crosstalk reduction to large void spaces in the image that represent the veins in the liver. Empty areas do not contribute to channel crosstalk and makes the correction inaccurate in the neighboring areas. This could be overcome, for example, with advanced methods able to detect the presence of veins in the image and adjust the crosstalk correction accordingly. All the images shown have been frame averaged 10 times and unwarped in *x* to correct for resonant scanning. The liver sample was cleared with SeeDB. Scale bar, 50 µm.

#### Crosstalk reduction

The factor *λ* that parametrizes the crosstalk reduction

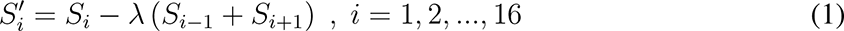

was optimized by minimizing the similarity between two adjacent strips. The variable *Si* represents the *i*-th strip of the FOV, which contains the fluorescence signal from the *i*-th anode of the PMT (Supplementary Fig. 7.a). To quantify the strip similarity, we used the structural similarity index (SSIM)^8^ on the 8th and 9th strips of the FOV. We plotted SSIM versus *λ* in cleared mouse liver tissue imaged close to the surface (Supplementary Fig. 7.b). We found an SSIM minimum at *λ∗* = 0.28 for the cell nuclei and *λ∗* = 0.35 for the plasma membranes, which we applied globally to the entire liver lobe. These values were obtained from averaging 5 measurements in random sample locations.

The inter-channel crosstalk, quantified through *λ∗* (the minimum of SSIM versus *λ*), increases slightly with the imaging depth (Supplementary Fig. 7.c). This is due to tissue-induced optical aberration that enlarges the fluorescence spots focusing onto the anodes of the PMT. Additionally, fluorescence photons scattered by deep tissue result in crosstalk between the multifocal detection cones at the sample plane. The dependency of the signal crosstalk on depth can be straightforwardly integrated into Supplementary Eq. 1, which was not implemented in this paper because of their minor effect.

**Supplementary Fig. 7:**
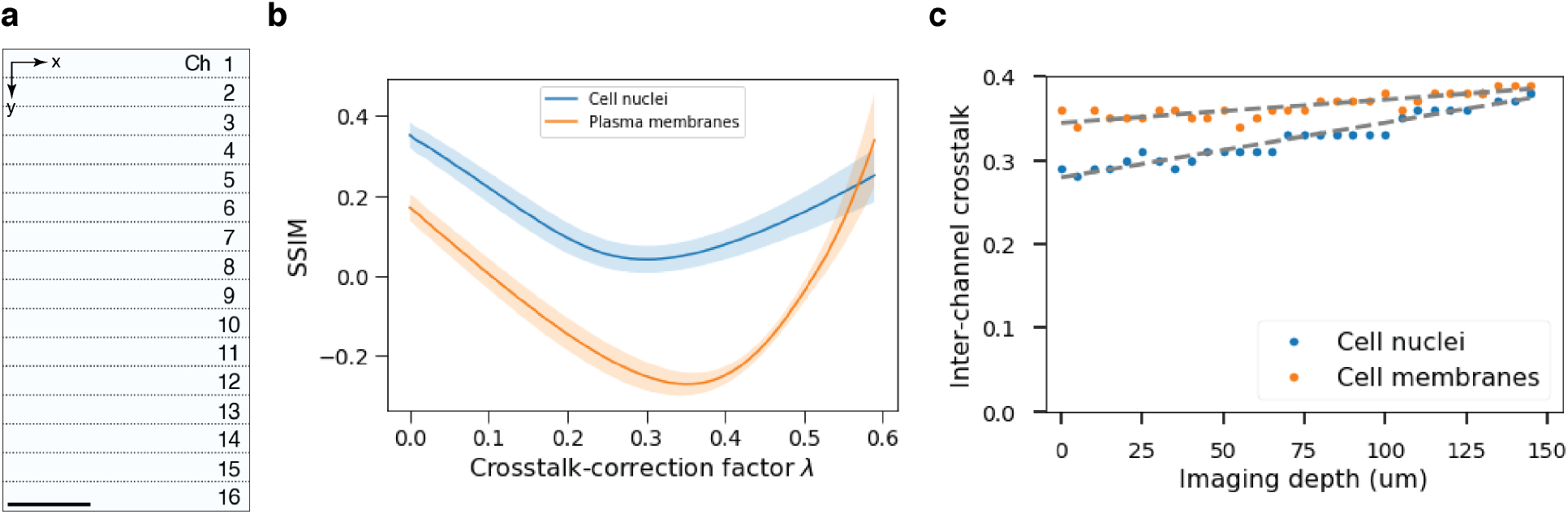
Crosstalk reduction. **a,** Field of view of 2pCOMB formed by 16 strips vertically concatenated. Each strip contains the fluorescence signal from each anode of the PMT. Scale bar, 50 µm. **b,** SSIM for different values of the crosstalk-correction factor *λ*. SSIM was computed between the 8th and 9th strips. The plot was obtained from averaging 5 measurements in random sample locations. The error bands show the standard deviation. The minimum was found at *λ∗ ∼* 0.28 for the cell nuclei and *λ∗ ∼* 0.35 for the plasma membranes in shallow tissue. **c,** Inter-channel crosstalk (*λ∗*) versus imaging depth. The signal crosstalk increased slightly with depth because of optical aberration and scattered photons in deep tissue.

#### Flatfield correction

The factor *α* that parametrizes the flatfield correction

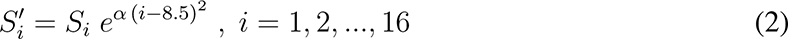

was determined by minimizing the Fourier component of the spatial frequency associated with the tile size in the *y*-axis. We found that *α* = 0.010 for the cell nuclei and *α* = 0.017 for the plasma membranes minimized the field unevenness. We applied these values of *α* to the entire liver lobe.

**Supplementary Fig. 8:**
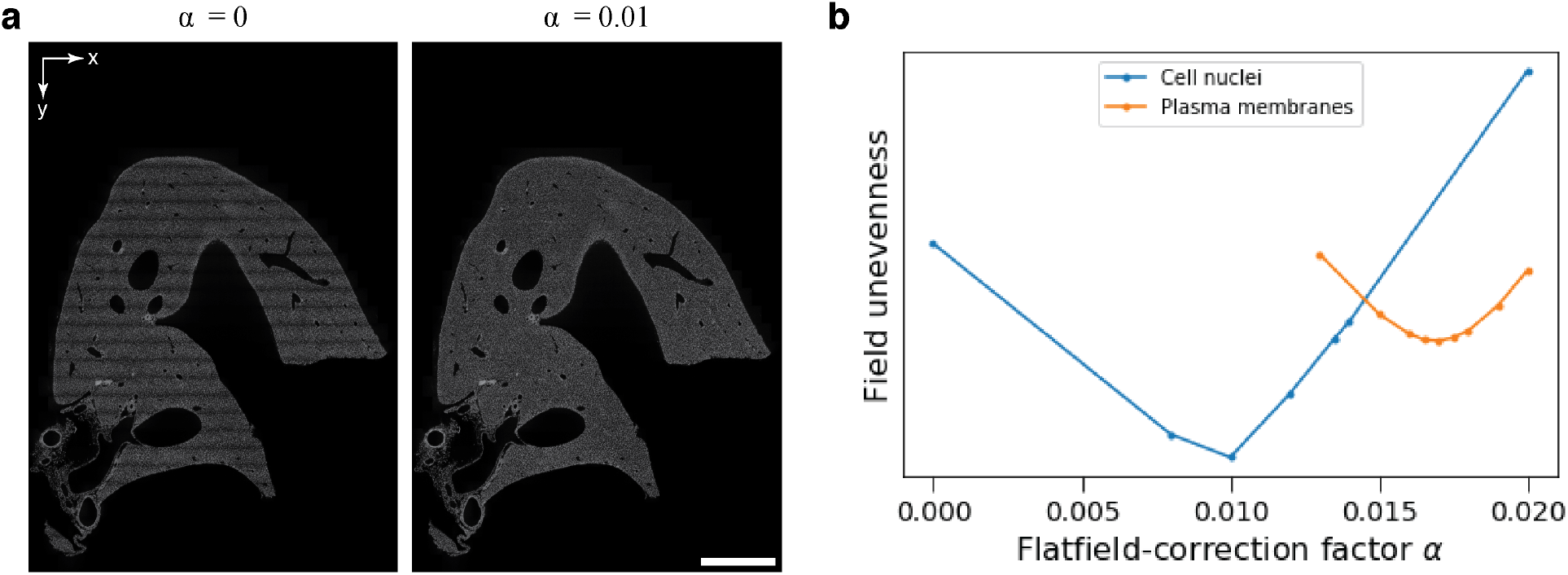
Flatfield correction. **a,** Cell nuclei across a *xy*-section after postprocessing for flatfield with *α* = 0 (left) or *α* = 0.01 (right). Scale bar, 1 mm. **b,** The unevenness of the image is quantified through Fourier analysis. The plot shows the Fourier component of the spatial frequency associated with the tile size in *y* versus the flatfield factor *α*.

Alternatively, we used a referential image from a uniform fluorescent slide, like the one shown in Fig. 2.a, to normalize the brightness of the FOV. The corrected images were comparable to those from applying Supplementary Eq. 1 for the plasma membranes (labeled with tdTomato) but worse for the cell nuclei (labeled with DAPI). We attributed the inferior performance in the latter case to the difference in refractive index between the fluorescent slide (1.51) and the liver tissue (1.49), which is has a greater effect on short wavelengths.

### 9 Deep imaging in mouse liver tissue

The images shown in Supplementary Fig. 6 were acquired near the surface of the tissue. To evaluate the performance of our multifocal microscope in deep tissues, we imaged the optically-cleared liver at a depth of 100 µm below the surface. We compared the postprocessed images with control measurements acquired with the same microscope under a conventional single-beam configuration (i.e., with the beam splitter bypassed). We observed that the cell nuclei and plasma membranes were still visible in deep tissues and that multibeam and single-beam scanning resulted in images of comparable quality (Supplementary Fig. 9). The images shown in Supplementary Fig. 9 correspond to those displayed in Fig. 2 in the main text.

**Supplementary Fig. 9:**
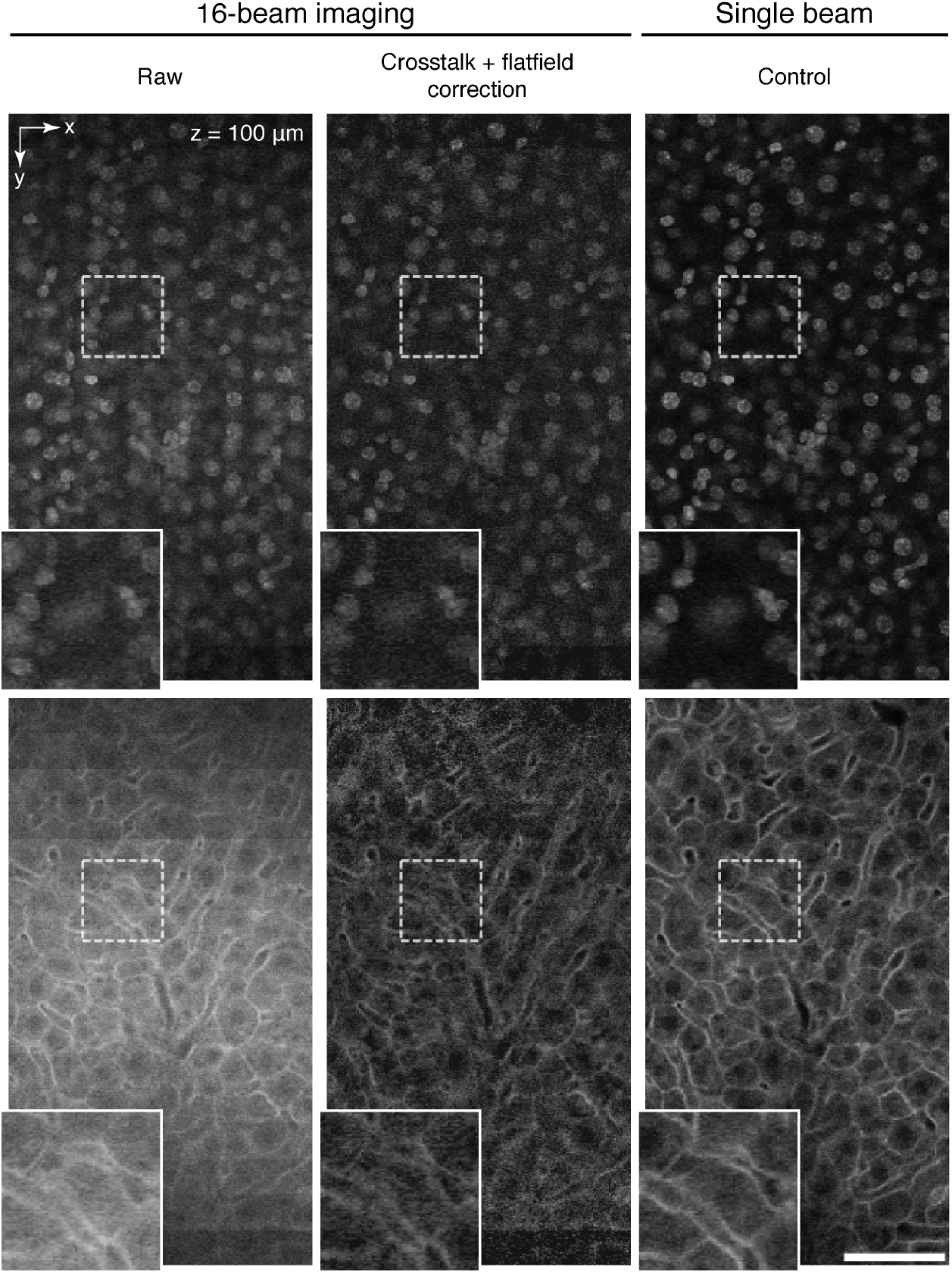
Deep imaging in mouse liver tissue. Cell nuclei (top row) and plasma membranes (bottom row) in mouse liver tissue imaged at a depth of 100 µm below the surface. The sample was cleared with SeeDB and imaged with 16 parallel beams via 2pCOMB with a frame average of 10. The control images (right column) show the same location in the tissue acquired with the microscope under a conventional single-beam configuration (i.e., with the beam splitter bypassed). The laser power was increased exponentially with depth. Post-acquisition correction was performed with *λ* = 0.33 and *α* = 0.010 for the nuclei and *λ* = 0.38 and *α* = 0.017 for the membranes All the images shown have been unwarped in *x* to correct for resonant scanning. Scale bar, 50 µm.

### 10 Image noise, frame averaging and deep-learning restoration

Images acquired via resonant raster-scanning (more commonly used for *in vivo* experiments for fast sampling) are shot-noise limited due to the low number of photons collected during the short pixel dwell time. In our experiments, we implemented a pixel dwell time of 162.5 ns, which allowed a maximum photocount of 13 per laser pulse at 80 MHz, as photocounting can only detect one photon per laser pulse. Nevertheless, the shot noise can be effectively reduced through frame averaging (Supplementary Fig. 10), Gaussian filtering, or deep-learning restoration^9^ (Supplementary Fig. 12), among other techniques.

**Supplementary Fig. 10:**
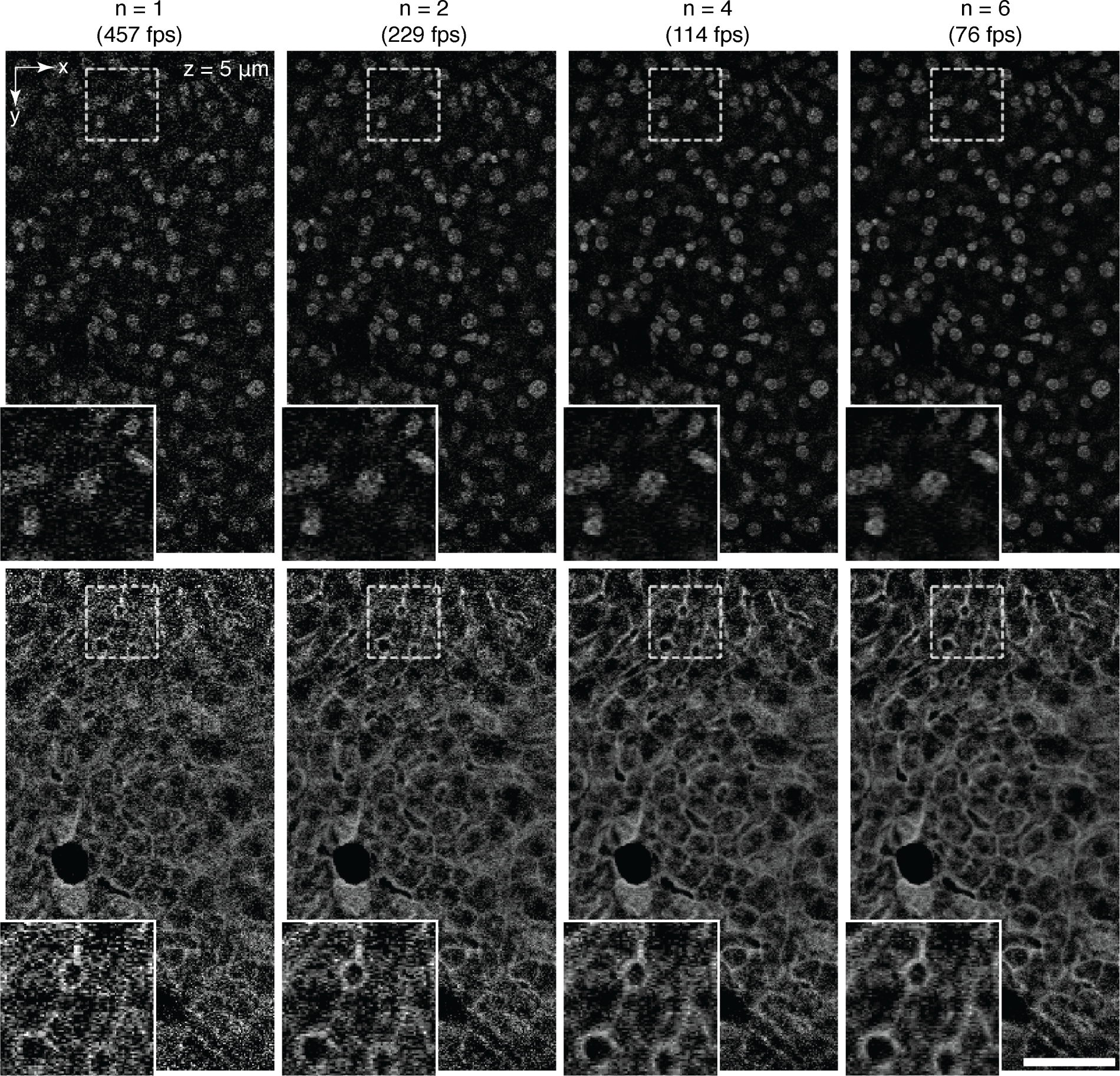
Image denoising via frame averaging. Cell nuclei (top row) and plasma membranes (bottom row) in mouse liver tissue acquired with 16 parallel beams via 2pCOMB with a frame average of n = 1, 2, 4, and 6. The corresponding acquisition rates are 457, 229, 114, and 76 frames per second, respectively. The liver sample was cleared with SeeDB. All the images shown have been postprocessed for field unwarping, crosstalk reduction, and flatfield correction The signal-to-noise ratio was lower on the top and bottom of the FOV than in the middle of the image due to the non-uniform laser intensities generated by the diffractive beam splitter (Online Methods). Scale bar, 50 µm.

**Supplementary Fig. 11:**
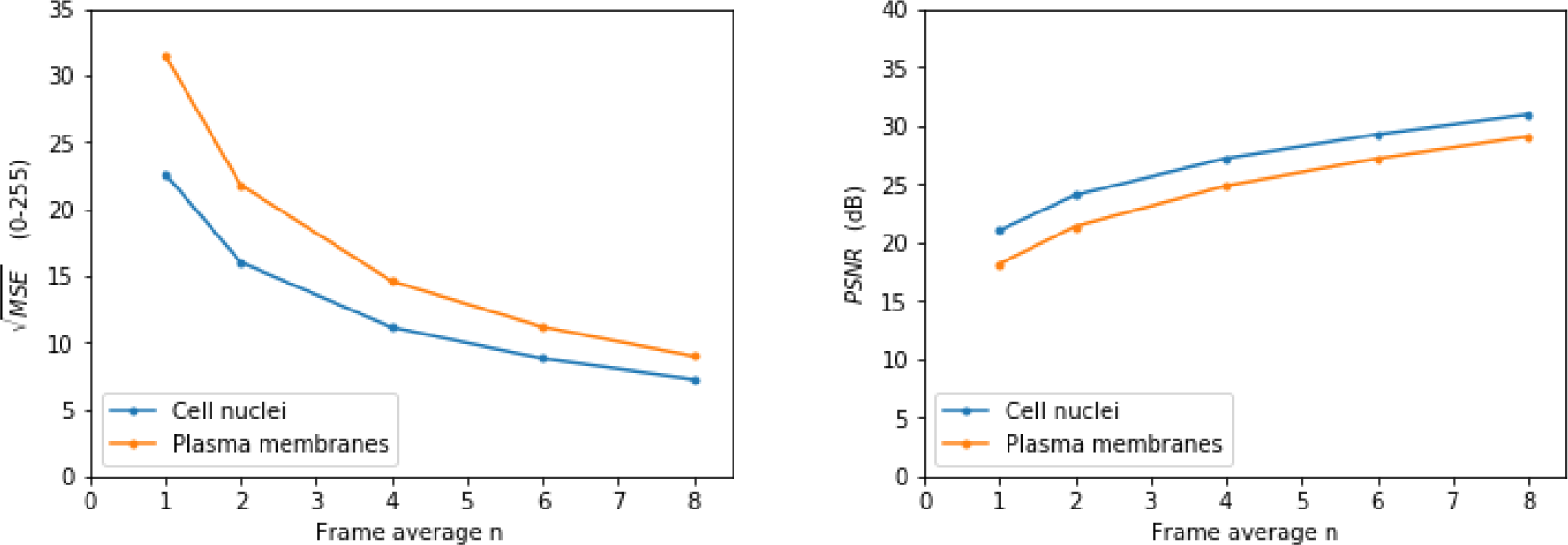
Noise level versus frame averaging. Square root of MSE (left) and PSNR (right) for the images shown in Supplementary Fig. 10.

The mean squared error (MSE) and peak signal-to-noise ratio (PSNR) plotted in Supplementary Fig. 11 are defined as

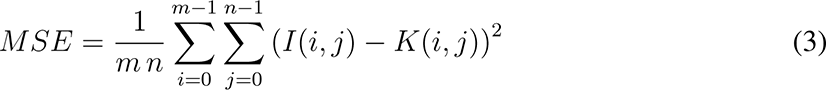

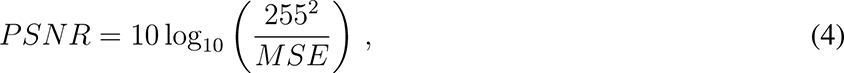

where *I* is a noise-free image and *K* is its noisy approximation. In practice, we used an image with a frame average of 10 for *I*.

**Supplementary Fig. 12:**
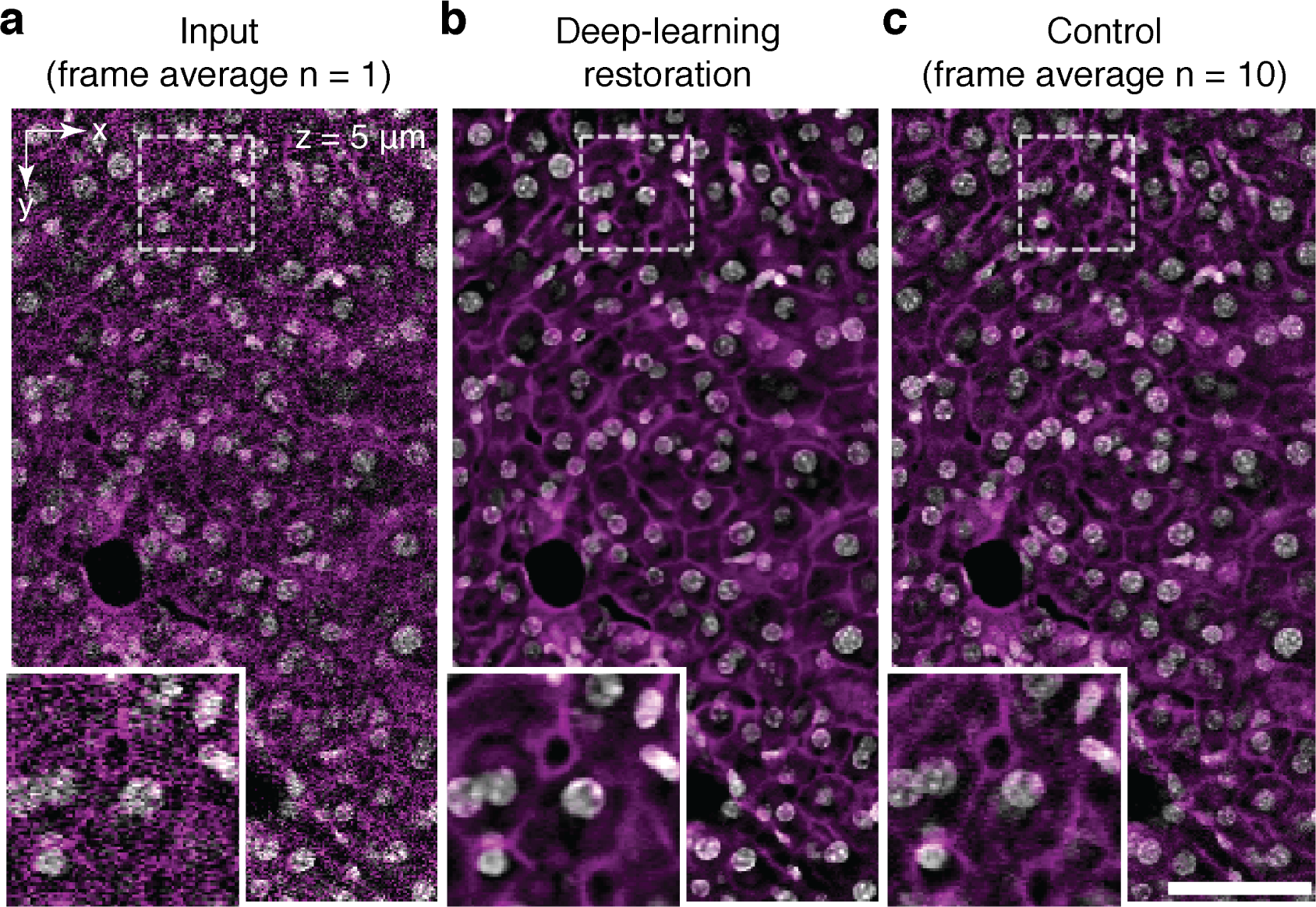
Image denoising via deep learning. Cell nuclei (white) and plasma membranes (magenta) in mouse liver tissue acquired with 16 parallel beams via 2pCOMB. Shot noise, common in resonant raster-scanning, can be denoised via deep-learning approaches. **a,** Image acquired at a multiplexed rate of 457 frames per second without frame averaging (n = 1). This image is identical to that of Supplementary Fig. 10. **b,** Denoised image after applying the deep learning method Content-aware Restoration (CARE)^9^. We trained the neural network with a collection of 50 pairs of 2D images (at low and high SNR) through the online implementation ZeroCostDL4Mic^10^. The training process took less than an hour with the default parameters, after which CARE could denoise a new 2D image in just a few milliseconds. **c,** Control image obtained from averaging 10 single-shot measurements. We performed CARE restoration on the raw microscopy data and then further postprocessed the images for field unwarping, crosstalk reduction, and flatfield correction. The liver sample was cleared with SeeDB. Scale bar, 50 µm.

### 11 Panoramic scanning of a mesoscopic sample

**Supplementary Fig. 13:**
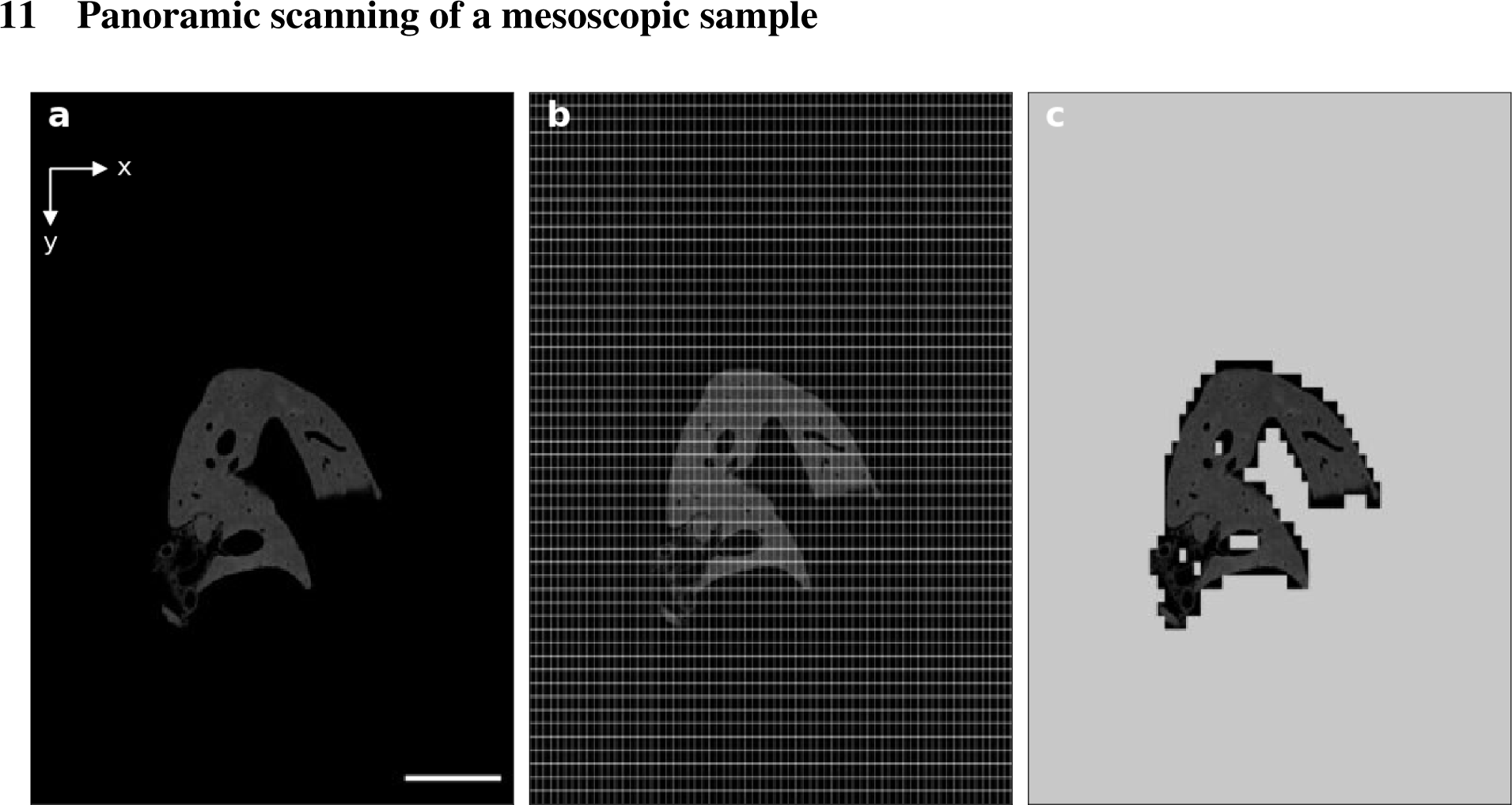
Panoramic scanning. **a,** Two-dimensional panoramic preview of the surface of the sample across a single *xy*-plane (~10 mm × 15 mm in *xy*). After cutting the sample with the vibratome, the location, extent, and shape of the new layers of tissue were previewed via panoramic scanning. **b,** A mosaic grid was then digitally overlaid on the image. The FOV of each tile was 150 µm × 280 µm in *xy*. The tile overlap was 10% in *x* and 15% in *y* (the overlap is imperceptible in the image). **c,** Only those tiles with an average pixel intensity above a certain threshold were subsequently volumetrically imaged via *z*-scanning. Scale bar, 2 mm.

### 12 Imaging parameters used for imaging the liver lobe

**Table 2:** Imaging parameters used for the liver lobe. The sample was imaged with 16 parallel beams via 2pCOMB. During acquisition, the sample was serially sectioned with an *in situ* vibratome. Each vibratome slice was *z*-scanned via mosaic acquisition (Supplementary Fig. 15.a).

|  |  |  |
| --- | --- | --- |
| Voxel | Size | $0.5 \times 0.5 \times 1 \mu\text{m}^3$ |
|  | Dwell time | 162.5 ns |
|  | Bit depth | 8 bits |
| z-stack | Volume | $280 \times 150 \times 100 \mu\text{m}^3$ |
| | Array size | $560 \times 300 \times 100$ voxels |
|  | Data size | 16.8 MB |
|  | Overlap in *x, y* | 10%, 15% |
|  | Acquisition time (frame average) | 438 ms (*n* = 2) |
|  | Time repositioning the *xy*-stages | ~250 ms |
|  | Time writing to disk (HDD) | ~100 ms |
| Vibratome slice | Thickness | 100 $\mu\text{m}$ |
| | Overlap in *z* | 50% (50 $\mu\text{m}$) |
| Full lobe | Dimensions | ~5 mm × 7 mm × 3 mm |
|  | Measured volume | 41.4 mm<sup>3</sup> |
|  | # vibratome slice | 52 |
|  | # stacks | 19,726 / label |
|  | Data size | 331 GB / label |
|  | Total acquisition time | ~11 hours / label |

For exceptionally long experiments (for example, in the case of multiple samples that are large and contain many fluorescence labels), the runtime could be further optimized by imaging without frame averaging, by reducing the stack overlap in *z*, or by parallelizing the acquisition sequence (for example, by saving a *z*-stack to disk and moving the *xy*-stages to the next grid position simultaneously).

### 13 Digital reconstruction of the liver lobe

#### Rigid stitching

The pipeline for volume reconstruction took the collection of post-acquisition-corrected *z*-stacks as input (~40,000 stacks, ~660 GB). The spatial coordinates of the stacks were stored in a configuration text file during imaging. A master script in Python called a series of scripts for (1) field unwarping, (2) crosstalk reduction, (3) flatfield correction, (4) pixel intensity normalization in *z*, (5) discarding the planes in each *z*-stack containing the tissue damaged by serial sectioning, (6) reformatting the configuration textfiles needed for stitching, (7) executing a Fiji macro for the plugin Grid/Collection Stitching that aligns and fuses the *z*-stacks within each vibratome slice independently, (8) executing a Fiji macro for the plugin BigStitcher that aligns and fuses the (independently fused) vibratome slices for reconstructing the full volume (Supplementary Fig. 14). Running the scripts from 1 to 7 took less than a day on a T630 Dell workstation with 24 cores and 382 GB of RAM. Fusing and exporting the whole volume with BigStitcher took ~50 hours at full resolution. When reconstructing partial volumes or downsampled versions of the full volume, we sometimes used Grid/Collection Stitching for fusing the vibratome slices because it performed significantly faster than BigStitcher. All the reconstruction steps can be carried out on computers with considerably less RAM and fewer cores than the workstation used.

The images and movies showing the 3D-rendered lobe were created with Imaris 9.5 installed on a standard personal computer. We downsampled the original voxel size of 0.5 µm × 0.5 µm × 1 µm in *xyz* to an isotropic spacing of 2 µm to be able to fit the whole lobe into RAM memory.

**Supplementary Fig. 14:**
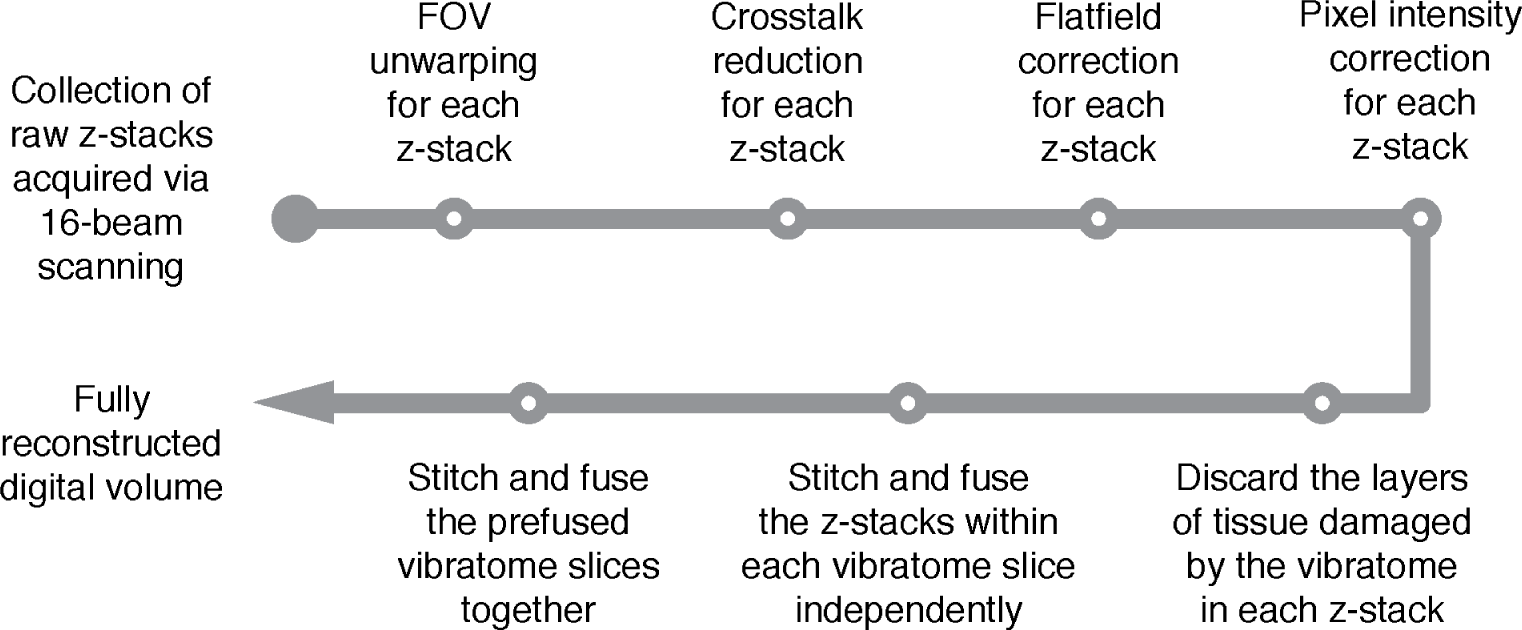
Pipeline used for reconstructing a mesoscopic digital volume from individual z-stacks.

#### Semi-rigid stitching

We implemented a semi-rigid stitching strategy in an attempt to improve the rigid alignment. Instead of directly fusing all the *z*-stacks in a vibratome slice, we created groups of 4 × 3 stacks that were aligned and fused (Supplementary Fig. 16.a). All the fused groups across all the vibratome slices were then jointly aligned with BigStitcher. This approach gives more degrees of freedom to the stitcher since groups within the same vibratome slice could move relative to each other. However, the alignment did not improve consistently. While the alignment improved in some regions (Supplementary Fig. 16.b), it deteriorated in others (Supplementary Fig. 16.c). We suspect that the strong correlations between groups in the same vibratome slice played a role in the lack of improvement. We, therefore, applied the simpler rigid approach described above to our data as it was sufficient for our goals of reconstruction and visualization. Future work could implement non-rigid approaches^11–13^.

**Supplementary Fig. 15:**
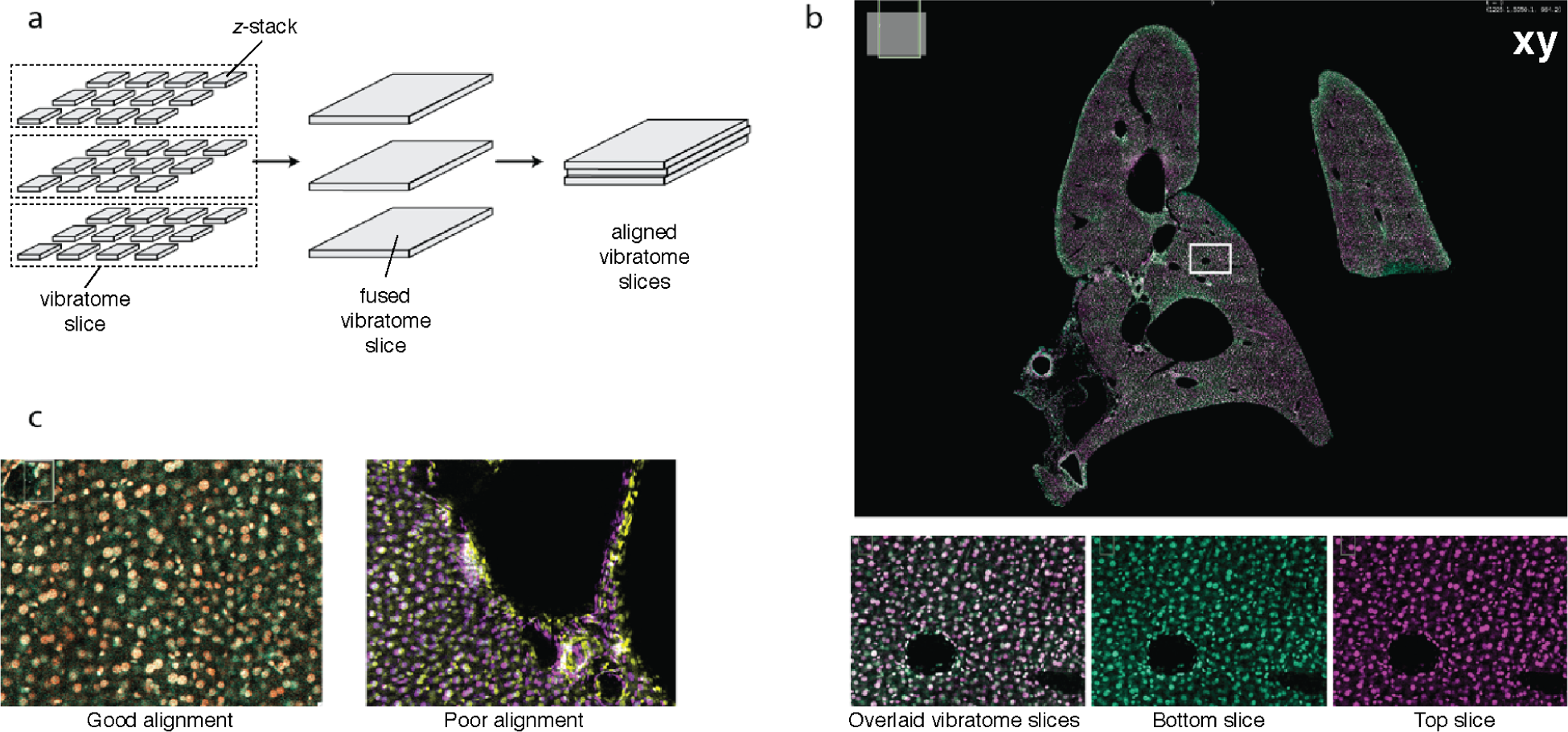
Rigid stitching of the liver lobe. **a,** Overview of the reconstruction pipeline. The individual stacks were first fused within each vibratome slice. The slices were then mutually aligned in *xyz* and fused. **b,** Overlaid *xy*-sections of 2 co-aligned vibratome slices. Only the cell nuclei are shown. The bottom slice is shown in green and the top slice in magenta. Inset: enlarged view of the selected area. The vibratome slices are shown overlaid (left) and individually (middle and right). The images are screenshots captured from BigStitcher. **c,** Example of alignment between neighboring vibratome slices showing good (left) and bad (right) performance.

**Supplementary Fig. 16:**
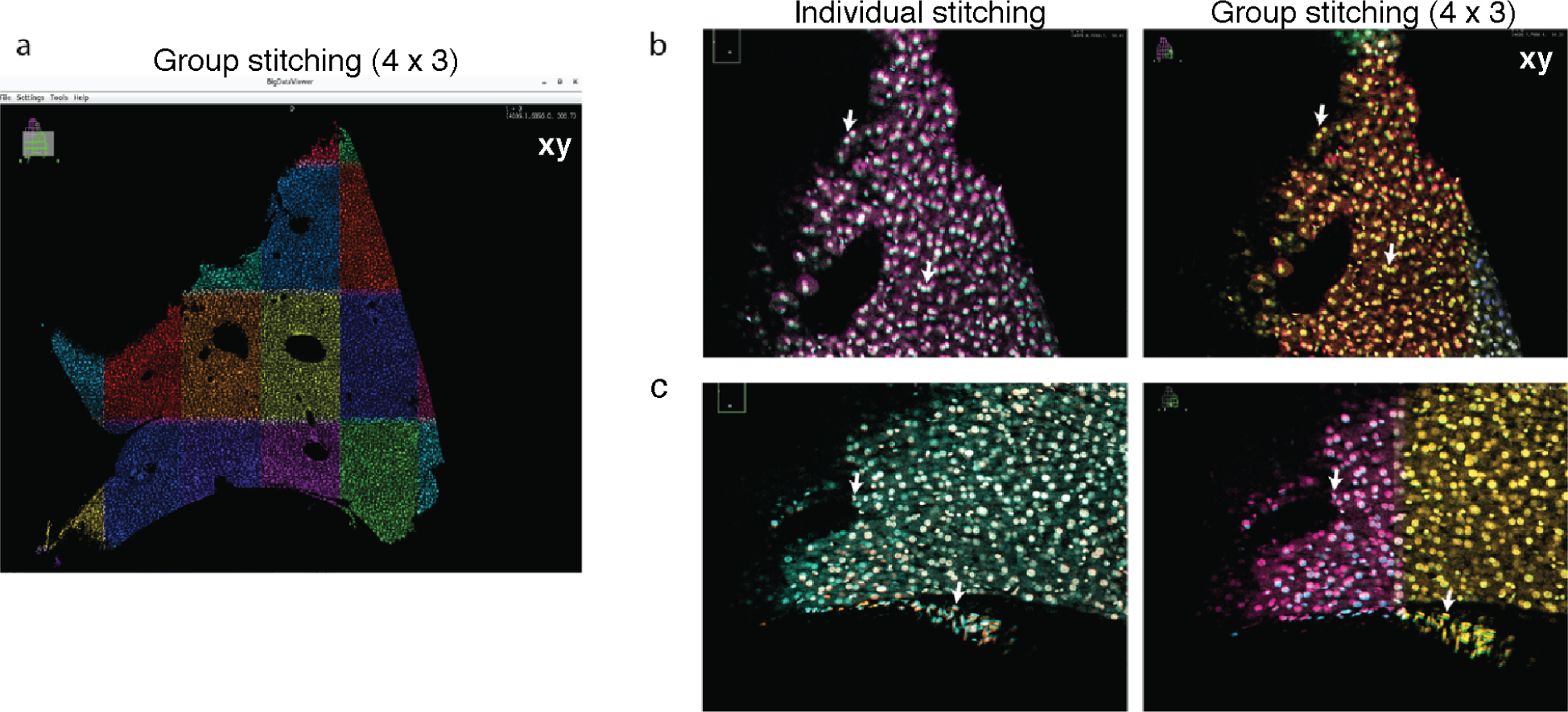
Semi-rigid stitching. **a,** Cross-section of a vibratome slice with the individual *z*-stacks fused into groups of 4 × 3 (shown in different colors). Each group is shown with a distinctive color. **b,** Cross-section of two contiguous vibratome slices co-aligned and overlaid in *z*. The alignment was performed via individual stitching (left column) or group stitching (right column). **c,** The same as in b at a different sample location. Group stitching led to inconclusive results. The quality of the alignment increased (b), decreased (c), or showed no effect. The arrows point to particular features to compare.

### 14 Description of the videos

Video 1: *xy* cross-section showing mouse liver tissue scanned with a single laser beam and 16 beams side by side. The cell nuclei are shown in gray and the plasma membranes in magenta. The voxel size is 0.5 µm × 0.5 µm × 1 µm in *xyz* and the frame average is 10. Postprocessing performed: field unwarping, crosstalk reduction, and flatfield correction.

Video 2: *xy* cross-section throughout a fully stitched and fused mouse liver lobe. The cell nuclei are shown in gray and the plasma membranes in magenta. The voxel size is 0.5 µm × 0.5 µm × 1 µm in *xyz* and the frame average is 2. The pipeline for post-acquisition processing is shown in Supplementary Fig. 14.

Video 3: A crop region of the full cross-section shown in Video 2.

Video 4: 3D rendering of the lobe shown in Video 2 with an isotropic voxel size of 2 µm. The cell nuclei are shown in gray and the plasma membranes in magenta.

Video 5: Zooming in on the 3D-rendered lobe shown in Video 4.

Video 6: Flying through the 3D-rendered lobe shown in Video 4.

Video 7: 3D rendering of a zoomed-in volume on the lobe shown in Video 4. The cell nuclei are shown in gray and the plasma membranes in magenta. The voxel size is 1 µm isotropic.

Video 8: Zooming in on the 3D-rendered volume shown in Video 7.

## Notes

### Competing Interest Statement

The authors have declared no competing interest.

